# Using network component analysis to study axenisation strategies for phototrophic eukaryotic microalgae

**DOI:** 10.1101/2025.03.07.641979

**Authors:** A. Iyer, M. Monissen, Q. Teo, O. Modin, R. Halim

**Affiliations:** Conway Institute of Biomolecular Research, University College Dublin, Belfield, Ireland; University College Dublin, School of Biosystems and Food Engineering, Belfield, Ireland; RWTH Aachen University, Templergraben 55, 52056 Aachen, Germany; Architecture and Civil Engineering, Chalmers University of Technology, Gothenburg, Sweden

**Keywords:** Axenisation, microalgae, eukaryote, phototrophs, single culture

## Abstract

**Background:** The axenisation of phototrophic eukaryotic microalgae has been studied for over a century, with antibiotics commonly employed to achieve axenic cultures. However, this approach often yields inconsistent outcomes and may contribute to the emergence of antibiotic-resistant microbes. A comprehensive review of microalgal species and the methods used to achieve axeny could provide insights into potentially effective workflows and identify gaps for future exploration.

**Methods:** Scholarly databases were systematically searched, supplemented by citation network analysis and AI-assisted tools, to collect studies on achieving axenic phototrophic eukaryotic microalgae cultures. Data about microalgal species, axenisation workflows, outcomes, and related factors (e.g., sampling locations, axenisation confirmation methods) were summarised. Network component analysis was used to identify clusters of commonly reported methods for diatoms, dinoflagellates, and green algae. A scoring framework was developed to assess the quality and reliability of evidence presented in the studies.

**Results:** Promising workflows circumventing the use of antibiotics appear to be filtration ↔ washing ↔ micropicking for diatoms, micropicking ↔ subculturing ↔ flow cytometry for dinoflagellates, and anoxy ↔ photosensitisation ↔ streak plating for green algae. Evidence from the literature indicates that a combination of microscopy (e.g., epifluorescence), cell counting (e.g., agar plating), and sequencing (16S and/or 18S) could enhance confidence in confirming axeny.

**Conclusion:** More systematic and high quality primary research is required to identify effective workflows for other microalgal divisions and fortify / contradict the ones proposed herein based on network component analysis.

## 1 Introduction

Axenicity “*describes the state of a culture in which only a single species, variety, or strain of organism is present and entirely free of all other contaminating organisms*” (Thain & Hickman, 2004). Popular culture banks host an impressive collection of microalgal species, but most of the offered type samples are xenic, i.e: they contain bacteria, amoebae, or fungi that grow in consort with microalgae in the natural habitat from where they are collected. Although characterising these commensal organisms provides important clues about the community where microalgae thrive, limiting the influence of confounders is crucial in experimental designs. For example, Deng et al. (2024) had to achieve and verify axenicity of their *Coscinodiscus radiatus* cultures to unambiguously assess the modulatory effects of the bacteria *Mameliella* sp. CS4 and *Marinobacter* sp. CS1 on microalgal aging and extracellular vesicle production. Robust investigation in microalgal research requires a supply of pure cultures, but despite a century of reportage on axenisation procedures no consensus or unified workflow has been described that can reliably generate pure cultures. Currently, the onus of achieving, maintaining and verifying axenicity of mother cultures used in experiments falls on the researchers rather than the supplier / culture collection. Consequently, interpreting the results from publications emanating from different research groups requires caution as the final outcome may have been significantly influenced by the co-cultured organisms.

Previously, Fernandez-Valenzuela et al. (2021) classified the methods based on their mode of action, and provided a narrative summary on their application to generate axenic microalgal cultures. Furthermore, they noted that decontamination required a tailored approach for each species, and highlighted the fragmented and inconsistent reportage of the methodology in the available literature. However, details on literature collection and analyses were not described in their review. Thus, the work presented herein attempts to systematically collect and collate all the available resources, extract the relevant information from the described methods, and perform network component analysis to understand overarching trends in axenisation workflows and address the research question: “*Which axenisation workflow(s) is/are best suited for a given microalgal division*.” The secondary aim was to identify axenisation workflows circumventing the use of antibiotics.

Network component analysis (NCA) is a useful technique for the comparison and study of different workflows as it can abstract the steps across multiple processes into a collection of nodes connected in a network, and help infer causal relationships within the network when direct measurements are not available. By understanding the sequence and combinations in which methods are applied, multiple pathways to effectively generate axenic cultures can be visualised and selected based on secondary considerations such as success rates of each method, or equipments available in a given facility.

Lastly, the results from this work are contextualised in real-world experiences and expectations by noting the comments from four culture banks on the challenges faced when attempting to generate and maintain axenic microalgal cultures.

## 2 Materials and methods

### 2.1 Search protocol

The search protocol used a hybrid method of structured and systematic querying using bibliometric databases, followed by citation network analyses, and artificial intelligence (AI) assisted services to discover and collect publications. The search term (algae AND (axen* OR purif*)) was queried in PubMed, Google Scholar, and Scopus bibliometric databases and filtered based on title and abstract using an awk script. The publications were then subjected to citation network analysis using the online tool Local Citation Network (https://localcitationnetwork.github.io/) and ResearchRabbit (www.researchrabbit.ai) to discover other publications based on input publications. The final list of recovered publications is available online at: https://www.researchrabbitapp.com/collection/public/0ZGR9O5YLQ. Publications in all languages were included and relevant information was extracted from the text using online translation tools. The working definition of microalgae for the purposes of this review was organisms that are single-cellular, phototrophic, eukaryotic, free-living, and capable of asexual reproduction. The exhaustive list of parameters, exclusion criteria and rationale is available in Supplementary Table S1.

### 2.2 Data items and synthesis

The following information was extracted from the literature and is provided in a master table as a .csv file (Supplementary Table S2):

1. Year of publication
2. Country of the research institution(s)
3. Microalgal species: The species, including the culture code / accession number if provided by the author.
4. Purification method(s): The method described by the publication to isolate microalgae or decrease the contaminant load. The order in which the methods are written represent the order in which they were applied in the described experimental procedure.
5. Screening method(s): Methods used to verify if the method used to achieve axenicity was successful. Alternatively, this also refers to methods to quantify the contaminant load and check the degree of success of the treatment.
6. Media used: The media used to grow the microalgae. In cases where the plates are used to test axenicity, the media composition of the agar plates was noted here as well.
7. Screening method(s): Methods used to verify axenicity. Alternatively, this also refers to methods to quantify the contaminant load.
8. Quality score: Refer to the subsection 2.3 below.

Data extraction procedures are detailed further in Supplementary Materials, Supplementary Text, Section 1.4. The various axenisation methods noted in the publications were classified either as Physical, Chemical, and Biological, where Physical methods involved the use of techniques such as colony picking to isolate a group of axenic cells from a Petri-dish for further culturing, or density gradient centrifugation to separate microalgae based on relative cellular densities. Chemical methods referred to the use of antibiotics or special compounds that inhibit the growth of unwanted microbes to achieve axenic microalgal cultures. Biological methods included methods that leveraged the unique abilities of the microalgae is used to isolate them from bacteria and fungi, for example, the use of phototaxis to isolate microalgae (Imai & Yamaguchi, 1994), or removing carbon source and depleting the nitrogen content of the medium, to give the microalgae a competitive advantage. Another example includes the addition of specific microbes that overgrow or inhibit the existing contaminating microbes, and then subsequently removing the introduced microbe using antibiotics or through physical methods.

### 2.3 Evidence qality

Using the approach described by Jadad et al. (1996), five questions were framed based upon which, the experimental quality of each publication describing the axenisation procedure was scored by AI, MM and QT. The final score was the rounded average of the three scores. The questions are noted below:

1. Does the publication report the source of the organism(s)? Yes: 1, No: 0 **Explanation:** The contaminating organism(s) may change based on the location from where the microalgae was sampled. In the case of purchase from culture collections, in-house processing protocols, long-term storage (such as cryopreservation), and handling may alter the composition or presence of other organisms cultured along with the microalgae.
2. Are technical details of the axenisation method explained to facilitate reproducibility? Yes: 1, Incompletely: 0, No: −1 **Explanation:** The question pertains to whether sufficient details of the chemicals or components used in the axenisation process is provided to easily procure or purchase them, or accurately construct, assemble or fabricate them.
3. Was growth medium clearly described? Yes: 1, Referenced: 0, No (either not mentioned, or only mentioned without proper reference): −1 **Explanation:** The growth medium works in synergy with the purification to support growth of the microalgae which may be stressed after any purification step. Corresponding changes made to the growth media would help ensure successful regeneration of the isolated cultures. Besides, numerous variants of media have been described without labelling or classifying them as an entirely different composition mix.
4. Was axeny verified after treatment? With multiple methods: 1, With only one method: 0, No: −1 **Explanation:** Veracity of the experimental purification method success validates treatment efficacy.
5. Were contaminating organisms identified? Yes: 1, No: 0 **Explanation:** A method’s efficacy may change depending in the nature of the species contaminating the microalgal culture. Characterisation or the identification of the contaminating microbes would help establish relevance to the purification method employed.

### 2.4 Data visualisation and availability

Data was processed using R (4.4.0) (R Core Team, 2024) and preliminary analyses was performed using the tidyverse package (Wickham et al., 2019). Phylogenetic trees from the species names were constructed using the R packages metacoder (Foster et al., 2017), ggtree (G. Yu et al., 2017), taxize (Chamberlain & Szöcs, 2013), and rentrez (Winter, 2017). Further network component analysis was performed using the iGraph (Csárdi et al., 2024) and tidygraph (Pendersen, 2024) packages. Subsequent visualisation of the network and the phylogenic tree was performed using the TikZ (Tanau, 2021) and pgfplots (Feuersänger, 2023) packages in LATEX (Lamport, 1994).

For the network analysis, two main outcomes were noted, firstly, the success rates for each method (number of times axenisation was successfully achieved / number of times the method was used), and secondly, the grouping / clustering of methods in a network for each microalgal division. To calculate success rates, only those methods that were reported in more than six species were included to remove over-representation of a single method or species within the division level analysis. Next, the clusters within each network were identified using “optimised modularity algorithms” (see documentation of iGraph) and the largest clique (methods commonly used together across the publications) was then obtained.

The codes used to process the information is presented in Supplementary_codes.pdf. The awk code used to filter additional irrelevant titles is provided in Supplementary Materials, Supplementary Text, Section 2. All data and resources necessary to reproduce the analyses is available in the OSF repository: https://osf.io/8cmk7/.

### 2.5 Contact with culture collections

The culture collections of CCAP, Bigelow, CSIRO and NIES were contacted to enquire about the number of unique microalgal species they house, and how many of them are available only in xenic forms, how many are available as axenic as well as xenic. In addition, a subjective question was asked on the challenges they face when trying to make some strains axenic. Their answers are quoted and added to the arguments made in the discussion section.

## 3 Results

### 3.1 Summarisation of data

The systematic search yielded 63 publications from which data was extracted and analysed as shown in Figure 1. Only four publications lacked any DOIs or similar permanent identifiers. Among the selected publications, 34 reported only one microalgae while the other 29 reported multiple microalgal species. Search query using databases yielded 52 relevant articles, and an additional 7 articles were recovered using citation network analysis, and another 4 using AI assisted methods. Thus lateral search methods could contribute to 17.7 % of the total recovered relevant publications. A summary of all the information collected from 63 publications is provided in Table 1. A consolidated master table with the greater details of each study is provided in Supplementary Table S2.

**Table 1:**
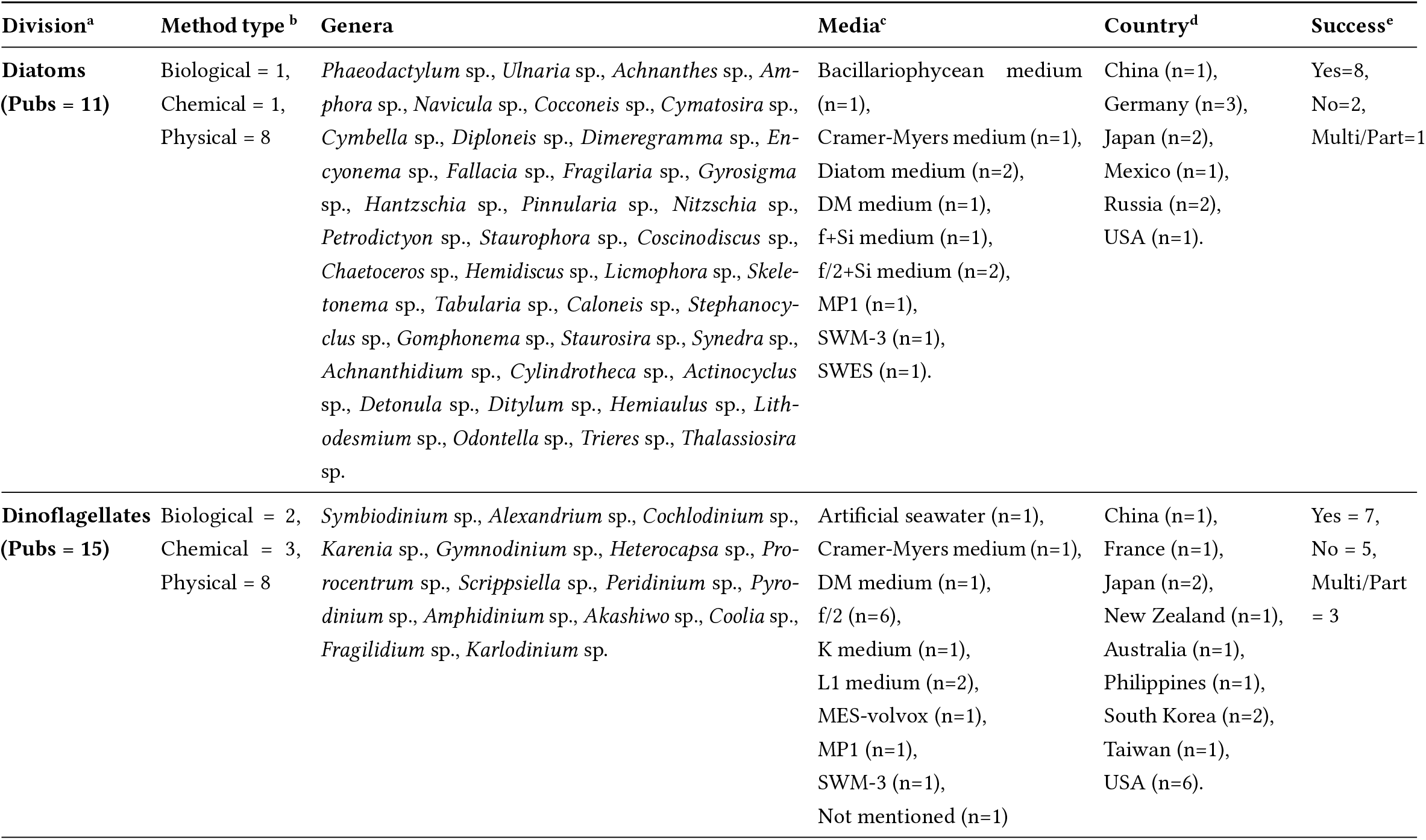
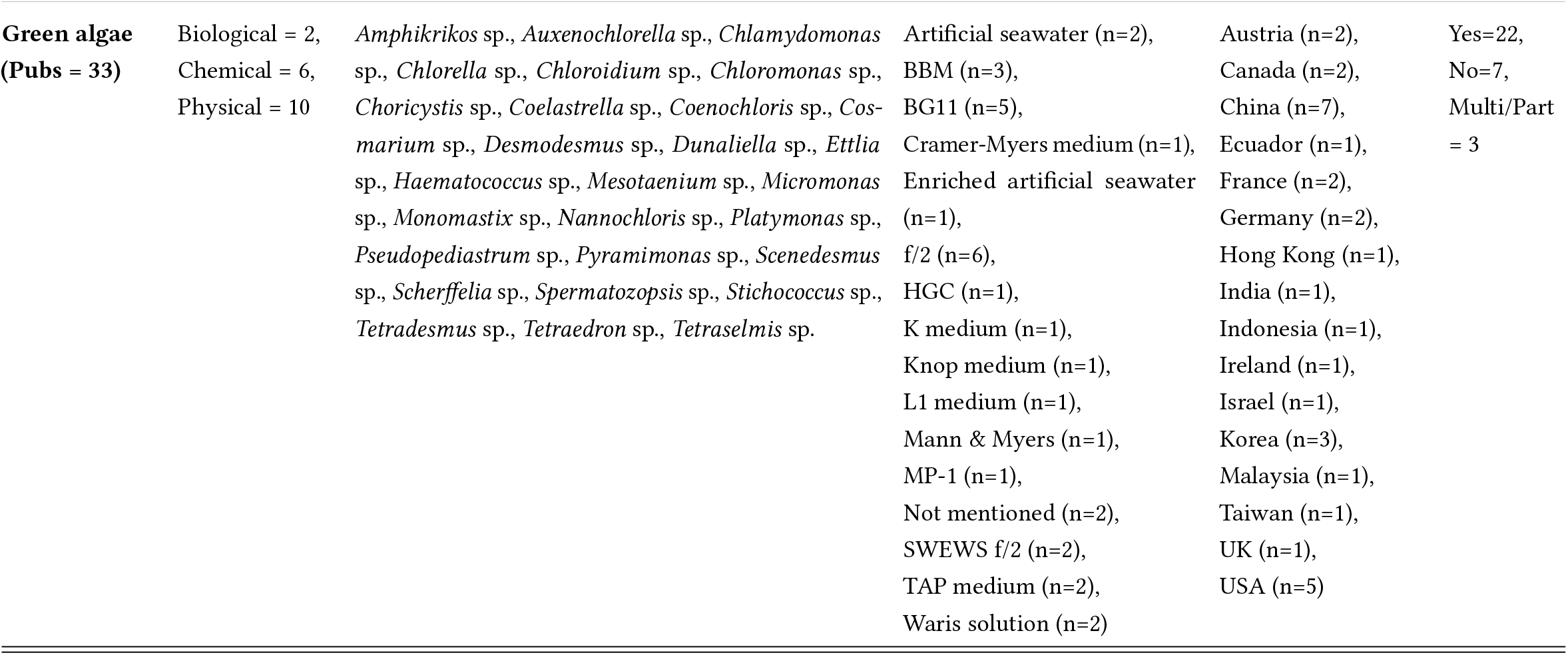
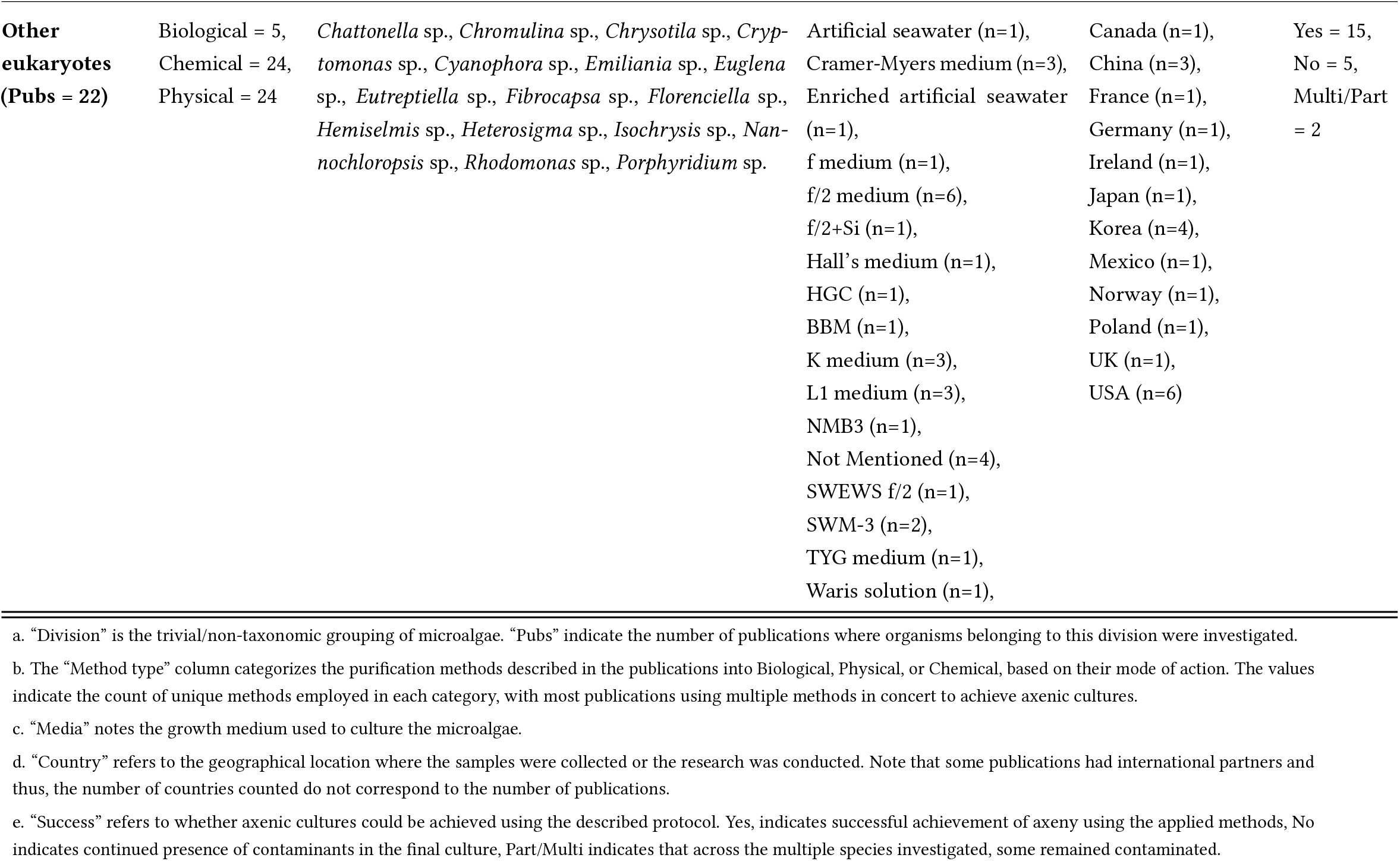
Summary table of the major data-points extracted from the recovered publications. Resolution of the information is at a publication level.

**Figure 1:**
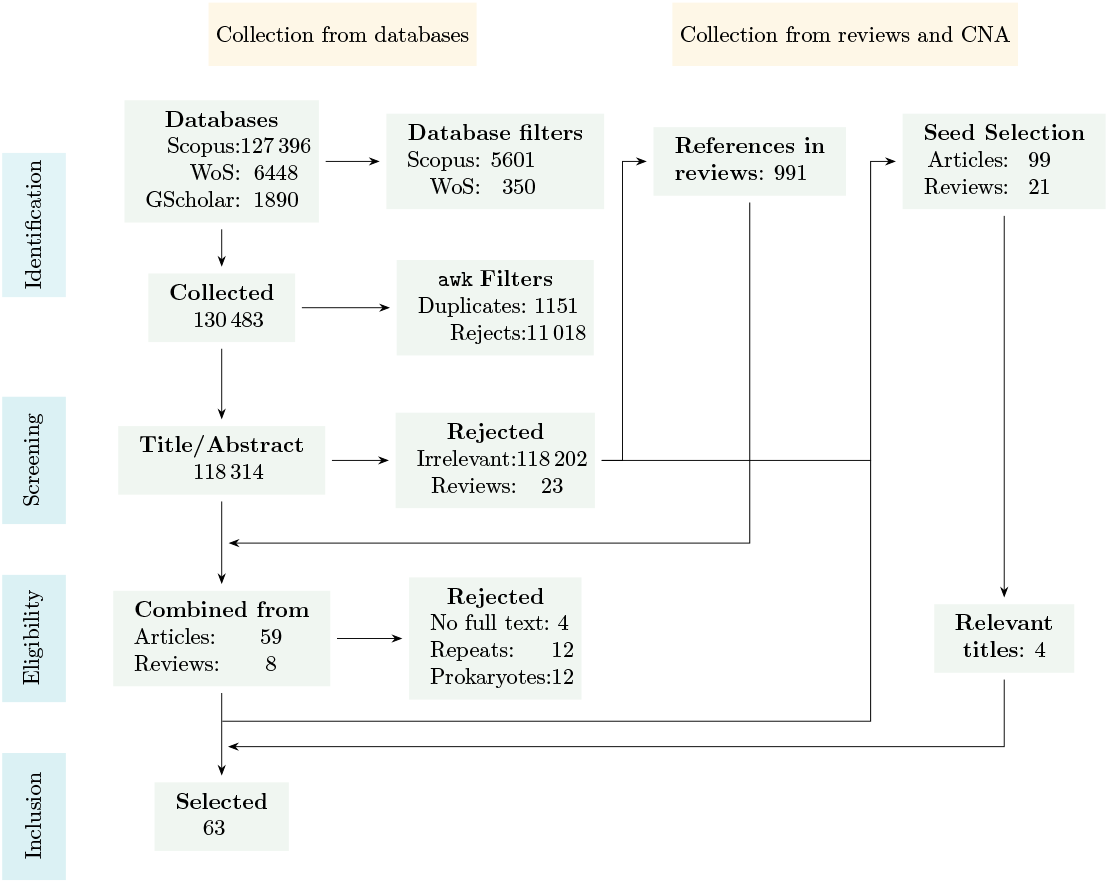
PRISMA-Sc flow schema of the search strategy and the recovered publications included herein. CNA = citation network analysis that was performed using the GitHub tool as well as ResearchRabbit to discover additional publications. Across the search workflow, 52 relevant articles using the traditional search method were recovered, with an additional 7 using citation network analysis, and another 4 using AI assisted methods. Thus lateral search methods could contribute to 17.7% of the total recovered relevant publications.

Across the publications, there were 24 unique axenisation methods that were broadly classified as “Biological” comprising of bacterial co-culturing (CoCu), selective predation (SePd), and phototaxis (PhoX), “Chemical” comprising of antibiotics (AntB), lysozyme (LysZ), hypochlorite (Chlo), surfactants (Dtrg), phenol (Phen), anoxy (Anox), dye-based photosensitisation (PhoS), and salt solution (Salt), and “Physical” comprising of centrifugation (Ctfg), streak plating (StPl), washing (Wash), micropicking (Mkpk), filtration (Fltr), density gradient (DenG), ultrasonication (UltS), subculturing (SubC), microfluidics (Mifl), french press (FrPr), serial dilution (SrDl), and adsorptive resin (Resn). A brief description of these methods are provided in Supplementary Text 1, Sections 1.1 to 1.3. The six most commonly used methods grouped according to the report quality and axenicity outcome is shown in Supplementary Figure S1 and the comprehensive counts of methods used for each species grouped according to according to axenicity outcome is provided in Supplementary Table S3.

### 3.2 Reported species and sampling locations

Across the literature, 174 unique species were identified and grouped into non-taxonomic divisions such as diatoms, dinoflagellates, etc. and further classified as freshwater, marine, or brackish species. Microalgae purified from soil were also found to grow in freshwater and were categorised accordingly. The recorded divisions included cryptomonads (n=4), diatoms (n=81), dinoflagellates (n=36), euglenoids (n=5), golden algae (n=1), eukaryotes (n=10), green algae (n=31), haptophytes (n=4), red algae (n=1), and green plants (n=1). Although diatoms comprised the largest number of reported species, they were generally subjected to only 11 of the total 24 methods. Conversely, all 24 methods were applied to the green algae division. Many of the reported organisms such as *Nannochloris* sp., and *Chlorella* sp., have significant commercial value and thus the investigation of newer methods to rapidly and effectively achieve axenic cultures is incentivized.

Only 19 publications documented the collection of samples from natural water sources (e.g., lakes, rivers, and coastlines) for the isolation and identification of microalgae. Furthermore, except for Aray-Andrade et al. (2018), who sampled along the Ecuadorian coastline, most reported sample collections from natural sources were confined to the Northern Hemisphere. Another 27 publications reported their experiments using cultures from culture banks (public or private), while 17 publications provided no information on where the microalgal samples were obtained.

The geographical sites where microalgal sampling were reported are shown in Figure 2, and the phylogenic trees of the reported species is presented in Figure 3.

**Figure 2:**
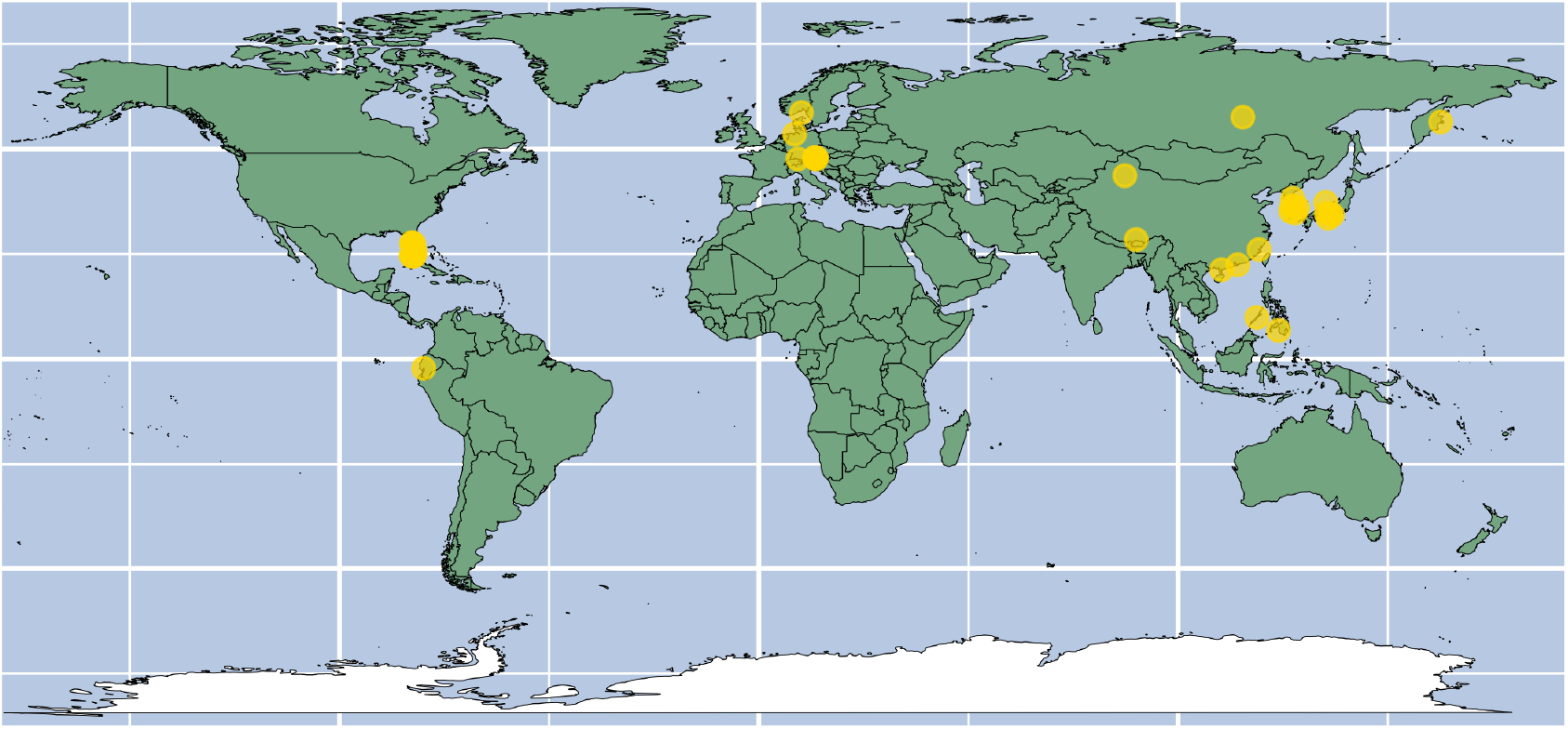
Geographic locations of microalgal samples collected from locations noted in 19 publications.

**Figure 3:**
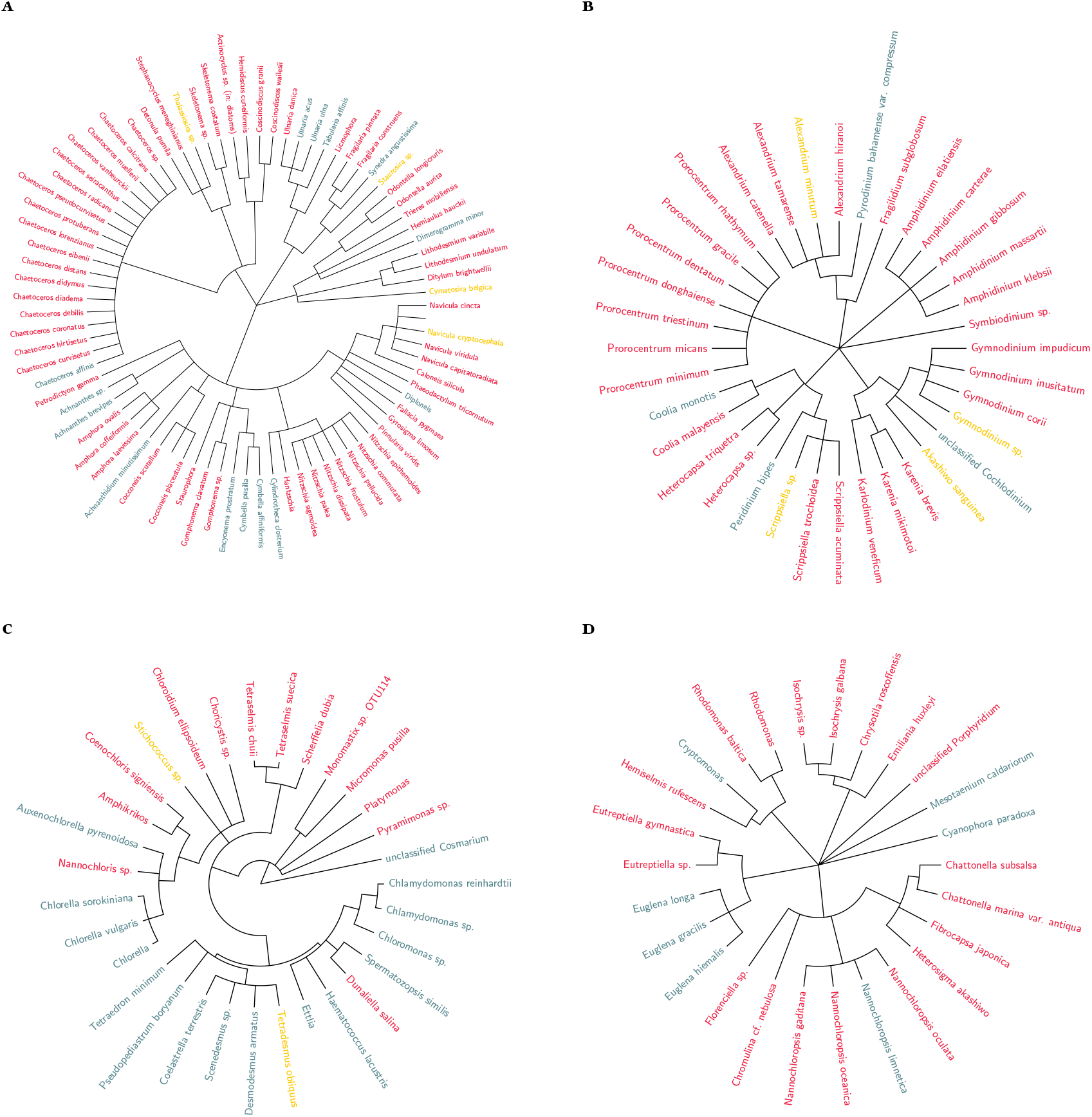
Phylogenetic tree of the species noted in the collected publications. Species names are coloured red for marine 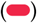, sunset-yellow for brackish 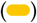, and steelblue 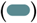 for freshwater species. (a) Taxonomic tree of 81 diatom species identified across 11 publications. (b) Taxonomic tree of 36 dinoflagellate species identified across 15 publications. (c) Taxonomic tree of 31 green algae species identified across 33 publications. (d) Taxonomic tree of 26 microalgal algae species identified across 22 publications.

### 3.3 Antibiotics – the most popular treatment

Antibiotics were by far the most preferred approach to generate axenic cultures as they are convenient to apply and the outcomes were generally rapid. Across the 63 publications, 42 reported the use of antibiotics spanning across 12 classes as shown in Table 2. In almost all cases, these antibiotics were applied as a cocktail and the combinations are shown in Figure 4. As the number of components in a cocktail increases, the range and potency to kill contaminants rose albeit, at the risk of affecting the viability of the microalgae.

**Table 2:** Antibiotic classes and concentrations used in microalgae axenisation workflows.

| Class | Diatoms | Dinoflagellates | Green algae | Others |
| --- | --- | --- | --- | --- |
| Aminoglycosides | 135 (10 to 600) | 160 (8 to 2000) | 600 (8 to 10000) | 600 (8 to 2000) |
| Carbapenems | 98 (98 to 98) | 10 (10 to 10) | 5 (5 to 5) | - |
| Fluoroquinolones | 5 (5 to 5) | 36.5 (23 to 50) | 13 (10 to 16) | - |
| Penicillins | 250 (50 to 600) | 100 (50 to 1000) | 600 (2.5 to 2000) | 100 (10 to 2000) |
| Tetracyclines | 10 (10 to 10) | 12 (1 to 12) | 50 (5 to 500) | 50 (12 to 500) |
| Cephalosporins | - | 60 (20 to 100) | 400 (10 to 500) | 30 (30 to 400) |
| Macrolides | - | 6 (6 to 6) | 54 (8 to 100) | 6 (6 to 6) |
| Polypeptides | - | 14 (12 to 16) | 3.3 (3.3 to 5000) | 16 (3.3 to 30) |
| Rifamycins | - | 10 (10 to 10) | - | 30 (25 to 30) |
| Nitrofurans | - | - | 10 (10 to 10) | 250 (250 to 250) |
| Antifungal Agents | - | 5 (5 to 15) | 20 (0.01 to 400) | 15 (1 to 15) |
| Others | 27 (27 to 27) | 125 (50 to 200) | 35 (5 to 60) | 20 (2.5 to 68) |

Values are expressed in $\text{mg L}^{-1}$ as median (minimum to maximum).

**Figure 4:**
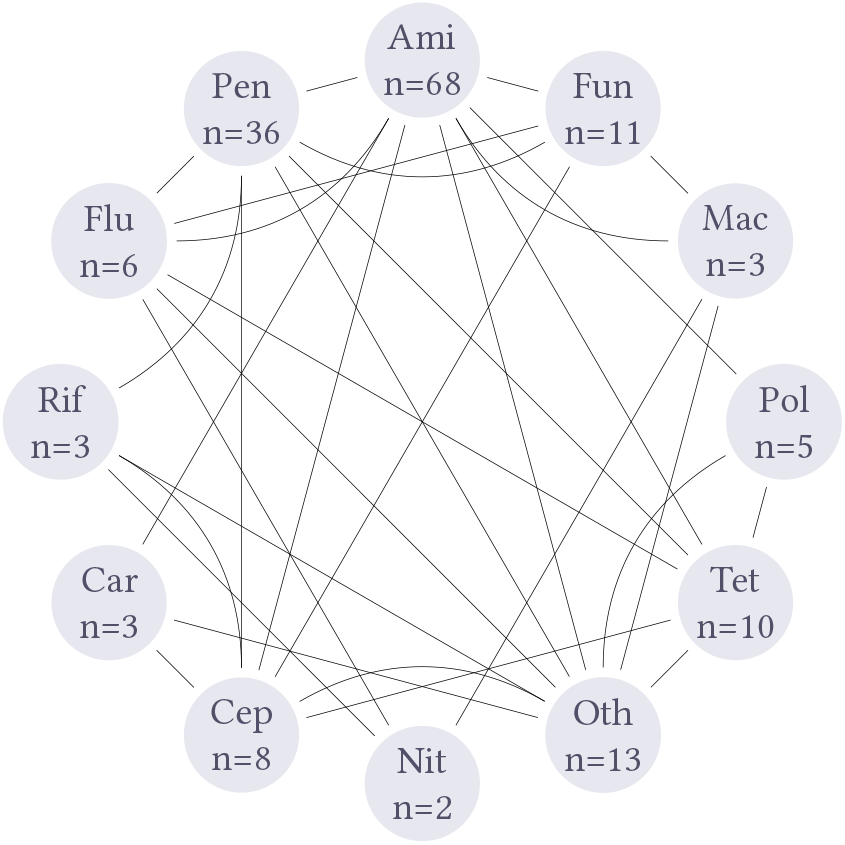
Antibiotic class combinations reported in the literature. Ami = Aminoglycosides, Fun = Antifungal agents, Mac = Macrolides, Pol = Polypeptides, Tet = Tetracyclines, Oth = Others, Nit = Nitrofurans, Cep = Cephalosporins, Car = Carbapenems, Rif = Rifamycins, Flu = Fluoroquinolones, Pen = Penicillins, n= number of publications reporting the use of the antibiotic class.

Diatoms were exposed to the fewest classes of antibiotics; predominantly Aminoglycosides and Penicillins, while dinoflagellates and green algae appeared to have been experimented upon with more classes of antibiotics. The three most commonly employed antibiotics included two aminoglycosides; namely, streptomycin (n=21; median conc. = 100 mg L^™1^), kanamycin (n=17; median conc. = 200 mg L^™1^) and one penicillin; namely, ampicillin (n=16; median conc. = 100 mg L^™1^). neomycin + kanamycin (n=7), streptomycin + penicillin-G (n=7) and streptomycin and neomycin (n=6) were the most common cocktail pairs employed in the publications. Supplementary Table S4 provides the name, class and concentrations of all reported antibiotics.

Antibiotics required dosing depending axenisation strategy. For example, Kan and Pan (2010) and Han et al. (2016) profess acute exposure of the culture to antibiotic cocktails to achieve axenicity, while Youn and Hur (2007) and Wang et al. (2016) employ antibiotics at moderate to low concentrations to achieve axenic cultures over multiple rounds of treatment to achieve and maintain axenicity.

### 3.4 Alternate axenisation pathways with network component analyses

Workflow networks for three of the most commonly reported divisions, namely, diatoms, dinoflagellates and green algae are provided in Figure 5a, 5b and 5c respectively. Workflow networks for microalgal families with low representation grouped under “others” is available in Supplementary Figure S2

**Figure 5:**
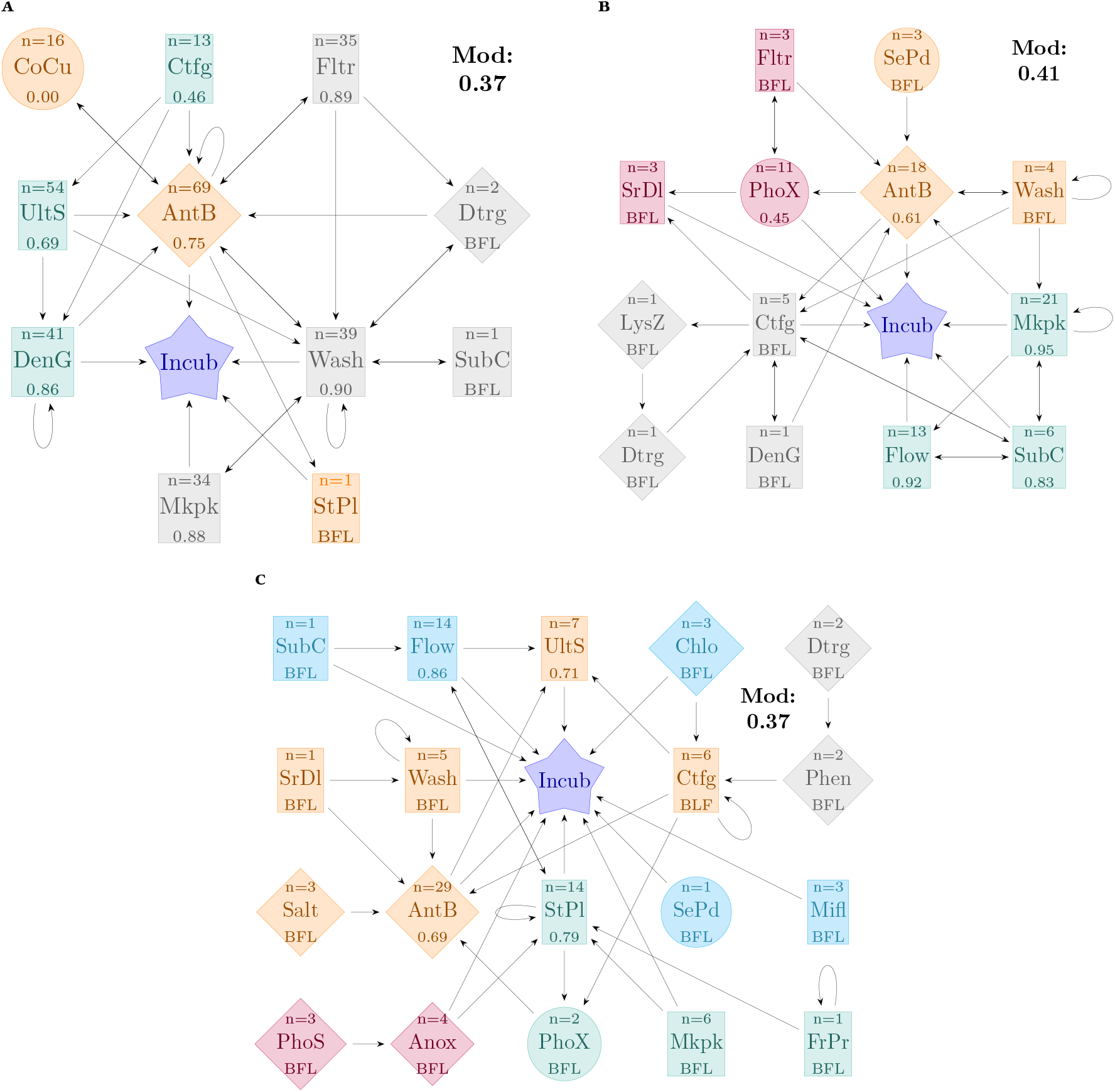
Clustering of the axenisation workflows. In each panel, methods belonging to the same cluster are assigned the same colour. Panel A shows the workflows incorporating the 11 methods used for 81 diatoms species in 11 publications. Panel B shows workflows incorporating 13 methods for 36 dinoflagellate species in 15 publications. Panel C shows the workflows incorporating 19 methods for 26 green algae species in 22 publications. Numbers in the network components represent success rates when number of reports is n>6; for n≤6 reports, success rates are below the filter limit (BFL) and not calculated. Node shapes denote method types: squares for “Physical”, circles for “Biological”, and diamonds for “Chemical”. Nodes of the same colour indicate they belong to the same cluster. The analysis is presented at the species level. Mod value represents the modularity value (range between 0= no clustering to 1 = perfect clustering) of the nodes in the network space.

Analyses included identifying clusters and cliques between the methods employed. “Clusters” represent the collection of nodes (here, methods) with similar network characteristics such as the edge density (number of connections to other nodes), and relative position and separation from the central node - Incubation (i.e: if they are directly connected to incubation or first connect to other nodes / methods before reaching incubation; see iGraph documentation by Csárdi et al. (2024) and more general works by Clauset et al. (2004) for deeper analysis). “Cliques” on the other hand comprise of adjacent nodes that are all directly interconnected (i.e: methods that are often used together in immediate succession) (Eppstein et al., 2010).

Network component analysis was applied to identify the largest clique of methods within a cluster so that methods with similar network properties could be used together. For example in Figure 5a, although filtration (Fltr), surfactants (Dtrg), washing (Wash), subculturing (SubC), and micropicking (Mkpk) clustered together (cluster 2, marked in grey) owing to similar network properties, the largest clique of methods was between Filtration ↔ Surfactants ↔ Washing. Details of the analyses and codes for Figure 5, are provided in Supplementary Codes (Code Chunk 6).

In the case of green algae (Figure 5a and Table 3), the overall network density was 0.11 (ratio of actual number of connections to the theoretical maximum number of connections) while in comparison diatoms have overall network density 0.21. Considering that 19 different methods have been reported on green algae across 22 publications, while only 11 methods were reported on diatoms across 11 publications, it appears that authors have been more experimental with the axenisation of green algae, with a disperse reportage on a wider method collection as opposed to diatoms where the method set is relatively limited and reported more frequently across the publications.

**Table 3:**
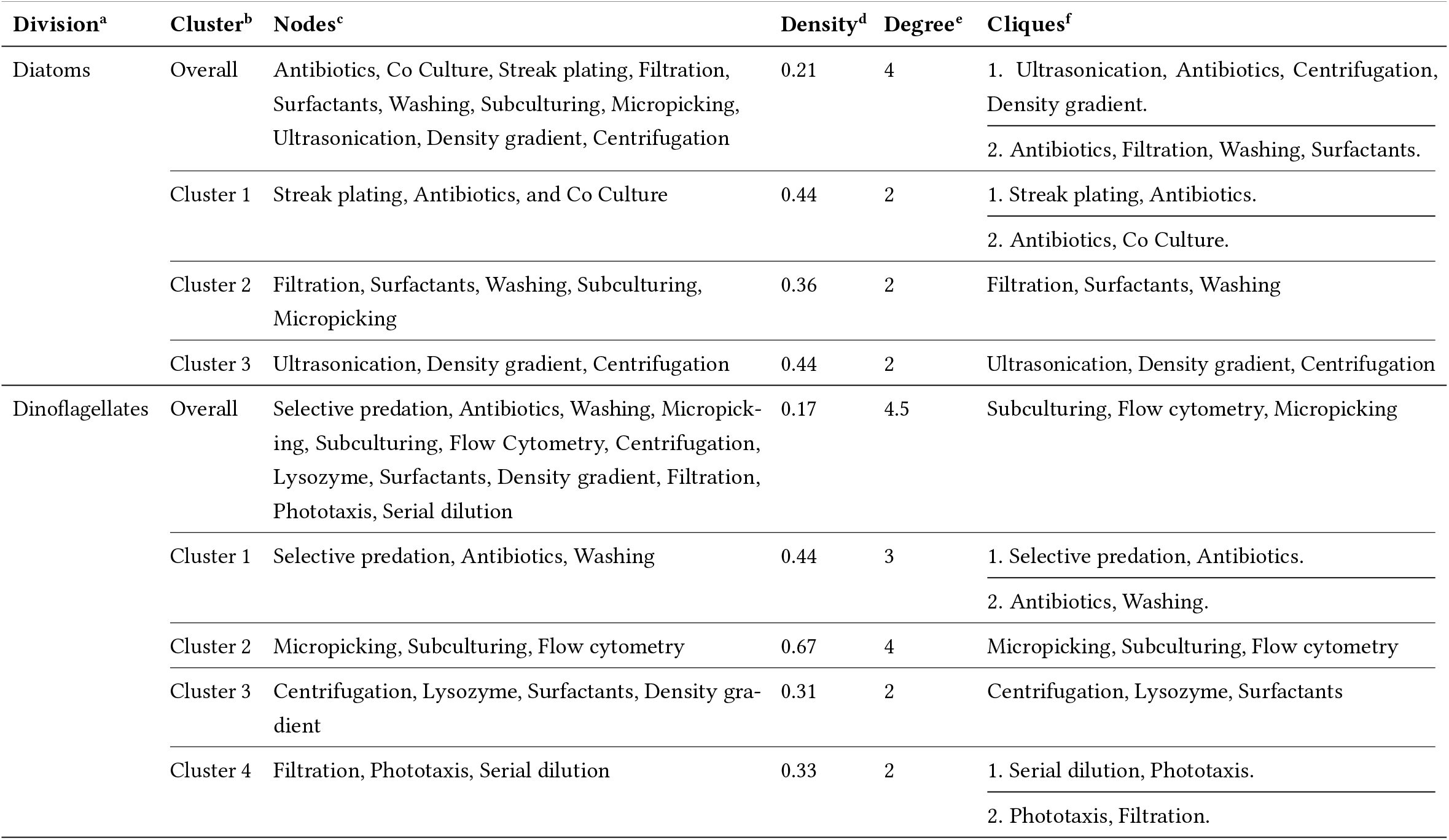
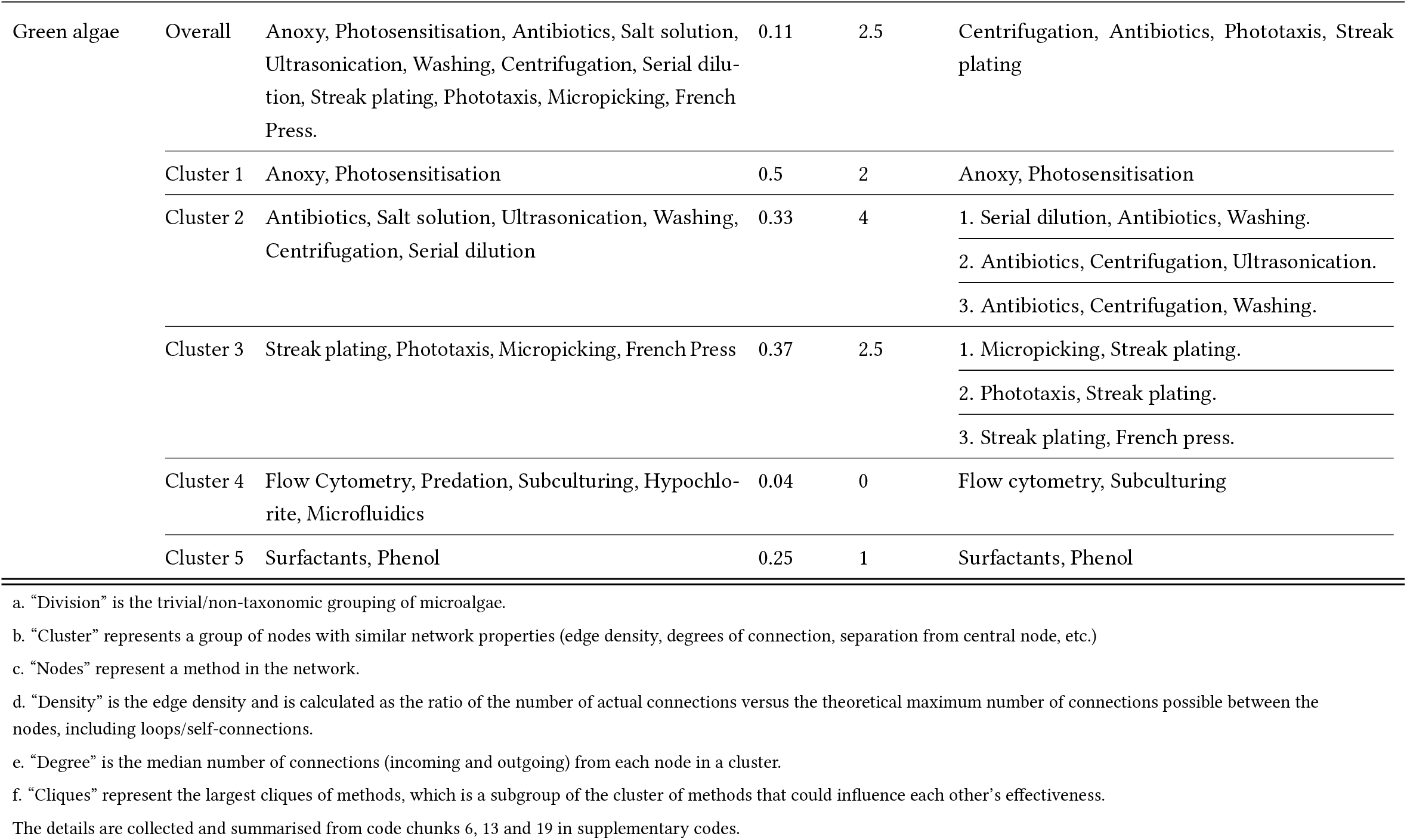
Network component parameters grouped by microalgal divisions.

| Division<sup>a</sup> | Cluster<sup>b</sup> | Nodes<sup>c</sup> | Density<sup>d</sup> | Degree<sup>e</sup> | Cliques<sup>f</sup> |
| --- | --- | --- | --- | --- | --- |
| Diatoms | Overall | Antibiotics, Co Culture, Streak plating, Filtration, Surfactants, Washing, Subculturing, Micropicking, Ultrasonication, Density gradient, Centrifugation | 0.21 | 4 | 1. Ultrasonication, Antibiotics, Centrifugation, Density gradient.<br>2. Antibiotics, Filtration, Washing, Surfactants. |
|  | Cluster 1 | Streak plating, Antibiotics, and Co Culture | 0.44 | 2 | 1. Streak plating, Antibiotics.<br>2. Antibiotics, Co Culture. |
|  | Cluster 2 | Filtration, Surfactants, Washing, Subculturing, Micropicking | 0.36 | 2 | Filtration, Surfactants, Washing |
|  | Cluster 3 | Ultrasonication, Density gradient, Centrifugation | 0.44 | 2 | Ultrasonication, Density gradient, Centrifugation |
| Dinoflagellates | Overall | Selective predation, Antibiotics, Washing, Micropicking, Subculturing, Flow Cytometry, Centrifugation, Lysozyme, Surfactants, Density gradient, Filtration, Phototaxis, Serial dilution | 0.17 | 4.5 | Subculturing, Flow cytometry, Micropicking |
|  | Cluster 1 | Selective predation, Antibiotics, Washing | 0.44 | 3 | 1. Selective predation, Antibiotics.<br>2. Antibiotics, Washing. |
|  | Cluster 2 | Micropicking, Subculturing, Flow cytometry | 0.67 | 4 | Micropicking, Subculturing, Flow cytometry |
|  | Cluster 3 | Centrifugation, Lysozyme, Surfactants, Density gradient | 0.31 | 2 | Centrifugation, Lysozyme, Surfactants |
|  | Cluster 4 | Filtration, Phototaxis, Serial dilution | 0.33 | 2 | 1. Serial dilution, Phototaxis.<br>2. Phototaxis, Filtration. |
| Green algae | Overall | Anoxy, Photosensitisation, Antibiotics, Salt solution, Ultrasonication, Washing, Centrifugation, Serial dilution, Streak plating, Phototaxis, Micropicking, French Press. | 0.11 | 2.5 | Centrifugation, Antibiotics, Phototaxis, Streak plating |
|  | Cluster 1 | Anoxy, Photosensitisation | 0.5 | 2 | Anoxy, Photosensitisation |
|  | Cluster 2 | Antibiotics, Salt solution, Ultrasonication, Washing, Centrifugation, Serial dilution | 0.33 | 4 | 1. Serial dilution, Antibiotics, Washing. |
|  |  |  |  |  | 2. Antibiotics, Centrifugation, Ultrasonication. |
|  |  |  |  |  | 3. Antibiotics, Centrifugation, Washing. |
|  | Cluster 3 | Streak plating, Phototaxis, Micropicking, French Press | 0.37 | 2.5 | 1. Micropicking, Streak plating. |
|  |  |  |  |  | 2. Phototaxis, Streak plating. |
|  |  |  |  |  | 3. Streak plating, French press. |
|  | Cluster 4 | Flow Cytometry, Predation, Subculturing, Hypochlorite, Microfluidics | 0.04 | 0 | Flow cytometry, Subculturing |
|  | Cluster 5 | Surfactants, Phenol | 0.25 | 1 | Surfactants, Phenol |

a. "Division" is the trivial/non-taxonomic grouping of microalgae.
b. "Cluster" represents a group of nodes with similar network properties (edge density, degrees of connection, separation from central node, etc.)
c. "Nodes" represent a method in the network.
d. "Density" is the edge density and is calculated as the ratio of the number of actual connections versus the theoretical maximum number of connections possible between the nodes, including loops/self-connections.
e. "Degree" is the median number of connections (incoming and outgoing) from each node in a cluster.
f. "Cliques" represent the largest cliques of methods, which is a subgroup of the cluster of methods that could influence each other's effectiveness.
The details are collected and summarised from code chunks 6, 13 and 19 in supplementary codes.

Details of the overall network, and the clusters and cliques for diatoms, dinoflagellates and green algae are summarised in Table 4. Identification of clusters and cliques for other eukaryotic microalgae (Supplementary Figure S2) was not performed as they did not represent any specific division / grouping.

**Table 4:**
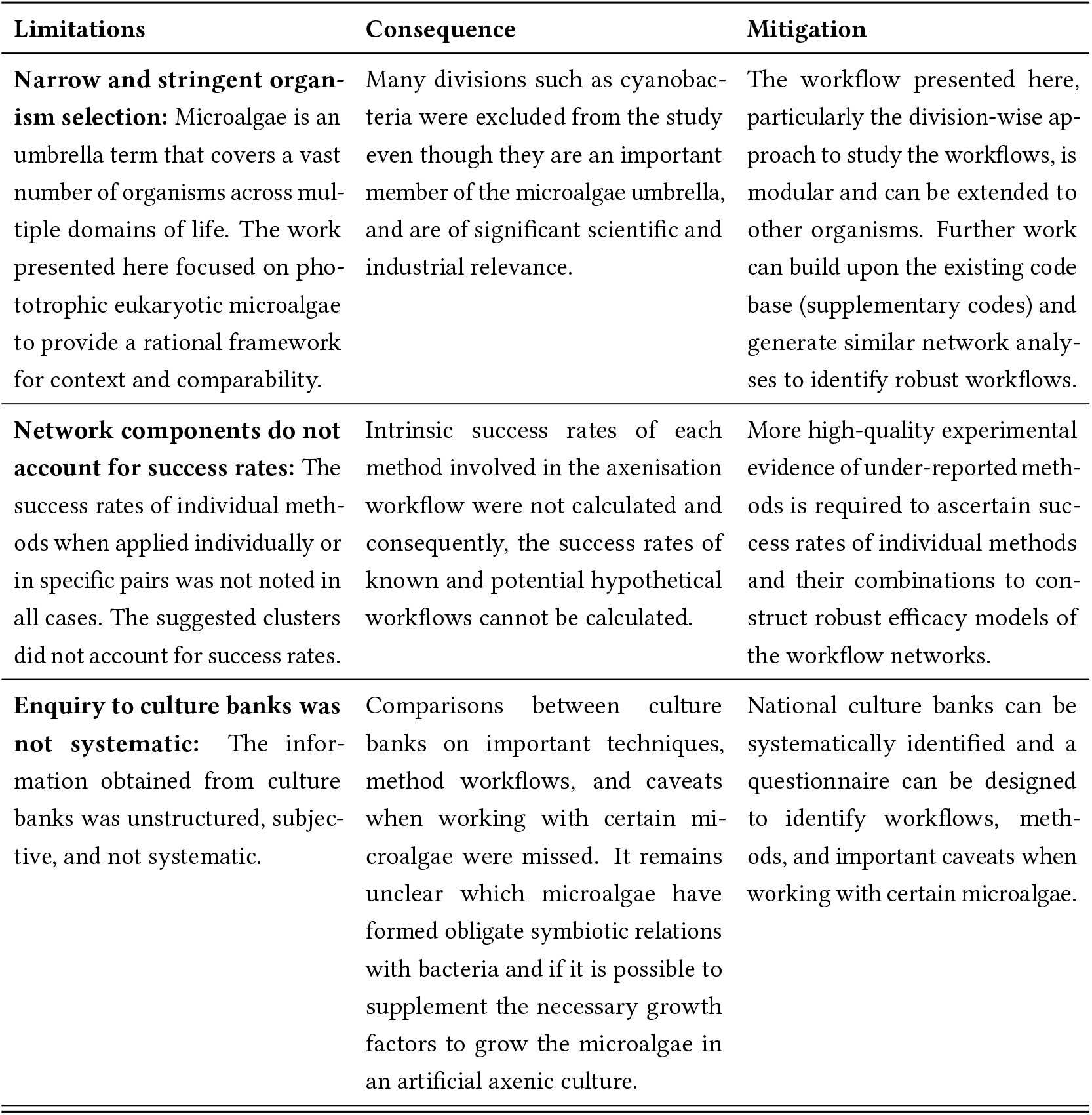
Limitations and caveats of the presented work.

| Limitations | Consequence | Mitigation |
| --- | --- | --- |
| **Narrow and stringent organism selection:** Microalgae is an umbrella term that covers a vast number of organisms across multiple domains of life. The work presented here focused on phototrophic eukaryotic microalgae to provide a rational framework for context and comparability. | Many divisions such as cyanobacteria were excluded from the study even though they are an important member of the microalgae umbrella, and are of significant scientific and industrial relevance. | The workflow presented here, particularly the division-wise approach to study the workflows, is modular and can be extended to other organisms. Further work can build upon the existing code base (supplementary codes) and generate similar network analyses to identify robust workflows. |
| **Network components do not account for success rates:** The success rates of individual methods when applied individually or in specific pairs was not noted in all cases. The suggested clusters did not account for success rates. | Intrinsic success rates of each method involved in the axenisation workflow were not calculated and consequently, the success rates of known and potential hypothetical workflows cannot be calculated. | More high-quality experimental evidence of under-reported methods is required to ascertain success rates of individual methods and their combinations to construct robust efficacy models of the workflow networks. |
| **Enquiry to culture banks was not systematic:** The information obtained from culture banks was unstructured, subjective, and not systematic. | Comparisons between culture banks on important techniques, method workflows, and caveats when working with certain microalgae were missed. It remains unclear which microalgae have formed obligate symbiotic relations with bacteria and if it is possible to supplement the necessary growth factors to grow the microalgae in an artificial axenic culture. | National culture banks can be systematically identified and a questionnaire can be designed to identify workflows, methods, and important caveats when working with certain microalgae. |

### 3.5 Appraising publications and confirming axenicity

A point-based appraisal mechanism was devised to assess publications describing experimental axenisation procedures for microalgae. Based on the questions noted in Section 2.3, about 7 out of 63 publications (11 %) had a score <0. The timeline of publications and the distribution of the publication quality within each time-period is provided in Supplementary Figure S3.

Poor method description (addressing Question 2) and lack of post treatment verification of axenicity (addressing Question 4) were the most common reasons for low scores. For example, across the 63 publications, six publications did not report a follow-up verification (consequently scoring −1 for Question 2), 23 publications reported the use of only one verification method (scoring a 0 for Question 2). Only 12 publications used a combination of two verification methods, while the remaining 22 employed three or more methods. A connection graph of the usage and combinations of verifications noted in literature is shown in Figure 6.

**Figure 6:**
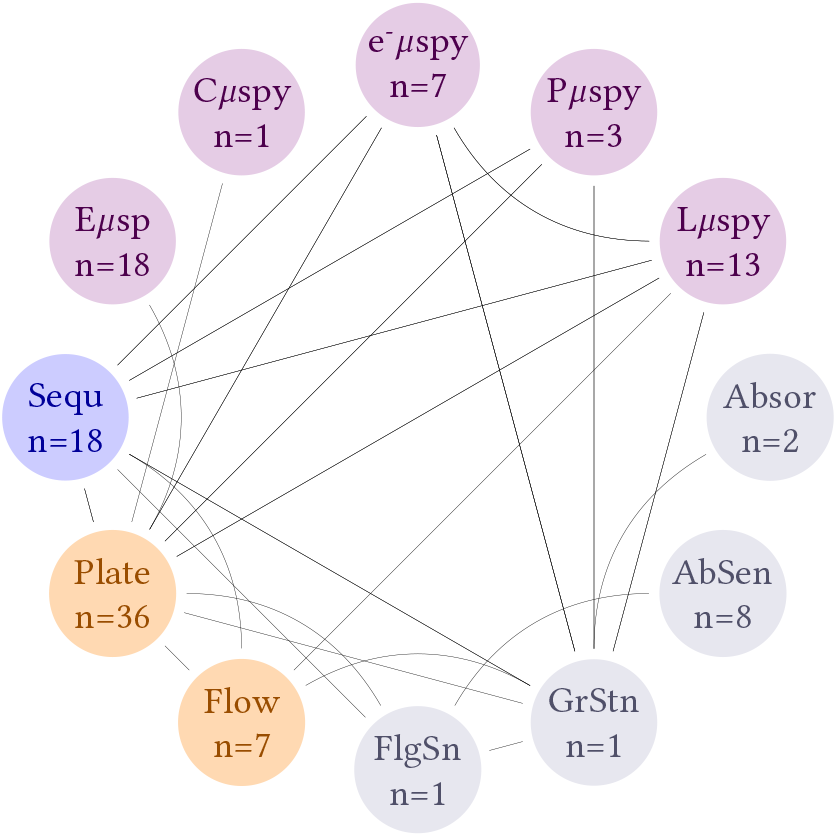
Methods reported in literature used either individually, or in combination to assess axeny in microalgal cultures. Resolution of information is at publication level. e-µspy = electron microscopy, Pµspy = Phase contrast microscopy, Lµspy = light microscopy, Absor = Absorbance, Flow = Flow cytometry, Eµspy = Epifluorescence microscopy, AbSen = Antibiotic sensitivity, GrStn = Gram staining, FlgSn = Flagella staining, Cµspy = Confocal microscopy, Plate = Plate counting, Sequ = 16S/18S sequencing. Methods marked in purple 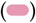 are microscopy-based methods. Methods marked in orange 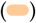 are cell counting methods. Methods marked in blue are sequencing methods 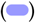. Methods marked in gray 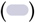 are physiology specific methods.

Across the publications, 13 distinct methods were reported to check axenicity of the final culture which could be grouped based on microscopy, cell/colony counts, sequencing, and physiology. Collectively, microscopy techniques that included confocal, electron, epifluorescence, phase contrast, and standard light microscopy, were the most popular approach to verify axenicity (n=42). Agar plate counting (n=36) was the next most common technique. Sequencing (both 16S/18S) methods were used in 18 publications, and lastly methods leveraging specific aspects of the microalgal physiology such as light absorbance, flagella staining, etc. were collectively reported in 12 publications.

## 4 Discussion

Axenisation of microalgal cultures remains a topic of significant research interest and can become necessary or undesirable depending on the research question being pursued. For example, Li et al. (2022) profile the bacterial consortium at each life stage of *Haematococcus pluvialis* and posit that the presence of bacteria enhances biomass production and growth-rate without any real threat of a culture crash, while B. S. Yu et al. (2022) contradict this by reviewing the strategies to purify the microalgae and note the risks posed to a cultivation batch under different nutrient conditions when contamination is unchecked and highlight the importance of maintaining axenic and sterile cultivation conditions to achieve optimal biomass and astaxanthin production.

On a wider note, there is a resurgence of interest in community interactions between microalgae and other organisms (Pushpakumara et al., 2023), either as interactions in non-associative biological mixtures in an experimental system (Grossart, 1999; Solomon et al., 2023), or indeed in symbiotic periphyton (Francoeur et al., 2003), and subsequently their emergent metabolic properties in relation to the wider ecology. Consequently, teasing out the individual contribution from a group effect, or attributing metabolic properties to an organism remains challenging (Fouilland et al., 2018). Although the publication by Dextro et al. (2024) focuses on cyanobacteria, the arguments noted in the review about the desirability (or lack thereof) of axenicity in microalgal cultures translate to phototrophic eukaryotic microalgal cultures as well, and their conclusions appear to suggest that for axenic cultures are suitable for genomic, proteomic, and industrial applications, while non-axenic cultures that preserve the natural bacterial consortium are necessary for fundamental research on understanding ecological dynamics between these microbes.

Many high quality and detailed reviews such as those by Fernandez-Valenzuela et al. (2021), McCracken (1969), Molina-Cárdenas et al. (2016), and Vu et al. (2018), have previously described axenisation procedures for microalgae. While these publications provide critical insights in applying the investigated methods, their approach was narrative and a top-down perspective of the various strategies adopted as a combination of axenisation methods has not been presented thus far. The approach adopted in this study was to collect methodologies described in publications and visualise them as “networks” of the methods connected to each other depending on the sequence in which they were applied that ultimately led to the final step of “incubation” where the culture is presumed to be axenic. This allows for methods to clusters within the network based on key characteristics such as edge density and relative node position. Edge density refers to the number of connections a node shares with others, influencing how tightly integrated a method is within the network. Another important factor is the relative position of a method relative to “Incubation” which is the final step in an axenisation workflow, and the central node in the network. This position determines whether a methodology is single step (i.e: only method directly connected to Incubation) or sequential (many methods used in sequence) before generating an axenic culture.

The methods described in the publications were grouped according to their mode of action into Physical, Chemical and Biological; with each presenting their characteristic advantages and shortcomings that need to be considered when designing an axenisation workflow. Physical methods require the most initial investment in the form of equipment as well as manual skill to successfully decontaminate the culture. By themselves physical methods such as density gradient, washing, or filtration do not generate axenic cultures, can contribute to the wider process to achieving them. Micropicking techniques require specialised skills to carefully separate microalgal cells from the contaminating organisms, without damaging cells. Flowcytometry on the other hand can automate the precise sorting process, but are generally ineffective against physical aggregates of microalgal cells and bacteria.

Chemical methods on the other hand are the most popular though the use of antibiotics. But as noted before, this has led to the wider problem of generating antibiotic resistant bacteria / fungi, while also adversely affecting the growth of the microalgal cells. In some cases, microalgae are known to metabolise the antibiotics and thus, antibiotic application may inadvertently become ineffective. Besides, chemical methods such as the use of detergents of oxidising agents such as chlorate, can significantly hamper the viability of the culture and incessant use can become mutagenic.

Lastly, biological methods have not been widely reported across the literature owing to the highly specific nature of their mode of action. Phototaxis is not a global microalgal ability, and is mostly restricted to certain dinoflagellates and green algae. Biological methods offer a more sustainable method of contaminant removal by leveraging physiological properties of the microalgae and selectively promoting their growth at the cost of the contaminants.

### 4.1 Clusters and cliqes from network component analysis

The network component analysis revealed variations in methodological approaches and interconnections across microalgal divisions, with key differences in network density, clustering, and reliance on specific methods. Workflows described to achieve axenic diatoms cultures comprising of 11 nodes (methods) exhibited a moderate network density of 0.21 (ratio of the number of connections to the theoretical maximum number of connections possible) and a median node degree of 4 (i,e: number of connections; incoming and outgoing, for each node), and an evenly distributed number of methods in each clusters. Workflows for dinoflagellates comprising of 13 nodes had a lower network density of 0.17, despite a higher median degree of 4.5, suggesting a broader range of methods applied, but with weaker interconnections. For example, methods such as phototaxis, or selective predation were used in limited combinations with other methods. The sparseness in the workflow network is demonstrated further in the methods described for green algae. With 19 nodes, the network density was only 0.11, and had a median degree of 2.5. This indicates that a large number of specialised methods were used, largely in isolation. For example, selective predation and microfluidics were used in isolation without using them in concert with other methods.

#### 4.1.1 Cluster characteristics

##### Diatoms

Two method-specific clusters of similar comprising of cliques with similar network densities of 0.44 emerged, namely Cluster 1 comprising of streak plating, antibiotics (success rate= 0.75), and co-culture (success rate= 0.0), and Cluster 3 comprising of ultrasonication (success rate= 0.69), density gradient (success rate= 0.86), and centrifugation (success rate= 0.46). Cluster 1 uses a mix of physical, chemical, and biological methods to achieve axenic cultures, while Cluster 3 comprises exclusively of physical methods and consequently, offers a workflow that circumvents the use of antibiotics.

##### Dinoflagellates

Cluster 2 comprised of a clique with the highest network density of 0.67, and exclusively involved physical methods of micropicking (success rate= 0.95), subculturing (success rate= 0.83), and flow cytometry (success rate= 0.92). Other clusters of interest were Cluster 3 and Cluster 4 that used methods leveraging the specificity of enzyme action (lysozyme) and motility (phototaxis), but had low network densities and insufficient number of publications to calculate success rates.

##### Green algae

Clustering with green algae was challenging as the method diversity was high, but the connections between them was weak owing to the lack of sufficient literature. For instance, Cluster 1 comprising of only anoxy and photosensitisation had the highest network density of 0.5, but none of the methods were sufficiently represented in the literature to calculate a success rate. In comparison, Cluster 2 comprising of a wider collection of methods such as antibiotics (success rate= 0.69), salt solution, ultrasonication (success rate= 0.71), washing, centrifugation, and serial dilution, and an overall network density of 0.33, had two methods with sufficient representation to calculate success rates, but involved the use of antibiotics.

##### Key comparisons and trends

Across the workflows described to achieve axenic microalgal cultures, clusters and cliques could be identified that avoided the use of antibiotics. For example, dense clusters for methods like micropicking, subculturing, and flow cytometry indicate that precision techniques could be used to achieve axenic dinoflagellate cultures. Similarly, with diatoms, filtration, washing and micropicking or ultrasonication and density gradient have been reported to attempt axenisation. In contrast, green algae does not show any preferred approach towards achieving axenisation, although there is preliminary evidence showing the use of anoxy and photosensitisation to be effective.

Across all divisions, antibiotics emerged as a common bridging method, appearing in high-degree nodes and multiple cliques. Mechanical methods like ultrasonication and centrifugation were consistently clustered in diatoms and green algae, but were less central for dinoflagellates, reflecting divisions-specific adaptations to their cell structures and ecological traits. Dinoflagellates had the highest reliance on precise isolation methods, likely due to their complex morphology and their own susceptibility to antibiotics and harsh chemical methods. Diatoms showed a balance between mechanical and chemical techniques, leveraging synergies within and across clusters. Green algae utilised a blend of unique (e.g., photosensitisation) and shared methods (e.g., antibiotics), indicating a diverse and disperse network of approaches.

### 4.2 Verification methods

Across the recovered literature, six publications did not properly verify the axenicity of their final culture, while eight publications only used one method to declare axeny. This is important as each verification method has inherent limitations that make it insufficient when used as the sole test. For example, plate counting is a well-established technique for detecting viable bacteria, as it requires microbial growth on a solid substrate. However, its limit of detection varies depending on the bacterial species and the growth conditions, typically ranging from >8 to 100 living cells / mL (Cappuccino & Sherman, 2008). Some bacteria may fail to grow on standard media but can persist in microalgal cultures, leading to false confirmation of axenicity. On the other hand, molecular approaches such as 16S rRNA gene sequencing are highly sensitive, with detection limits as low as ~10 cells/mL (Brandt & Albertsen, 2018). However, while sequencing can confirm the presence of bacterial DNA, it cannot differentiate between live bacteria and lysed cells. This means that sequencing alone may overestimate contamination levels by detecting residual DNA from non-viable cells. Microscopy methods, such as epifluorescence or differential interference contrast microscopy, provides direct visualization of potential contaminants but suffers inherent subjectivity in detecting the type and level of contaminants. The accuracy of this method depends on the experience of the observer and prior knowledge of expected contaminants in a given culture. Moreover, microscopy may fail to distinguish bacterial cells if they are present at very low densities or embedded within algal sheaths. Flow cytometry can be used as an alternative identify single cells as well as differentiate viability (using the Propidium Iodide or 7-aminoactinomycin D). However, the size disparity between microalgae and bacteria makes it challenging to accurately identify “singlets”, especially when dealing with closely associated bacteria-microalgae aggregates. For example, diatoms with tightly bound bacteria within the silica frustule may be detected as a singlet. Additionally, while live/dead staining can be incorporated into flow cytometry protocols to assess bacterial viability, this approach may still fail to detect bacteria in intimate symbiotic relationships with microalgae.

Given these limitations, a combination of complimentary testing methods is necessary to deduce axenicity with high confidence. Plate counting provides confirmation of viable bacteria, sequencing ensures highly sensitive detection, and microscopy serves as a cross-validation tool for detecting bacteria that might be overlooked due to primer biases in PCR or aggregation with microalgae. Flow cytometry could serve as a complementary tool, particularly if live/dead staining is included, but its efficacy depends on the strength of microalgal-bacterial associations. Based on the method combinations shown in Figure 6, a robust approach could involve using plate counting (or flow cytometry if live/dead staining is implemented) alongside sequencing and microscopy to maximize detection sensitivity while minimizing false negatives.

### 4.3 Perspective from microalgal culture collection

Culture collections are critical towards curating and maintaining microalgal cultures and was identified as a prime-source to gain insight from the expertise and knowledge drawn from regular practical work. A query was made to these culture collections to enquire about their practices to achieve axeny, and the tips and important considerations when maintaining axeny in working culture. The number of phototrophic eukaryotic microalgae along with the number of axenic species held in four major culture banks is provided in Supplementary Table S5. Upon enquiry, personnel from CSIRO stated that axeny was achieved through the use of antibiotics, while NIES exclusively used washing and micro-picking, and do not use antibiotics. The respondent from Bigelow did not mention any specific method, but acknowledged that the process can be labour intensive. All the culture collection respondents emphasised on the use of good sterile lab techniques to ensure axeny with the respondent from Bigelow noting that

> “Maintaining axenic cultures is not difficult if proper working conditions are maintained. We do all work in a flow hood and practice sterile technique.”

The emphasis of the importance of considering the role of commensal/phycospheric microbes was also universally echoed, with the respondent from CCAP stating,

> “… however we don’t try to make other axenic (unless necessary for a particular project) as many strains don’t like to grow well as axenic cultures. “

, and from CSIRO,

> “We find many do not grow if axenic, possibly due to the need for vitamins etc produced by bacteria. Streptomycins and G-penicillin etc can be used - some species will not tolerate them but most will.”

The respondent from NIES stated,

> “One of the challenges to maintain axenic strains is that not a few axenic strains grow poorly. This is likely due to the removal of benefit from coexisting bacteria, for example.”[sic]

It is unclear how many of the microalgae in the culture collection are obligate symbionts, and if there has been a systematic attempt to achieve and maintain axenic cultures. The responses from the culture collection institutes suggest that there is currently insufficient interest in the researchers requesting the cultures to generate and maintain axeny; likely ascribed to common understanding that beneficial and stable symbiosis between microalgae and pychospheric bacteria is needed to achieve high biomass productivity. The lack of interest can be due to the heavy initial investment and time/labour required for axenisation, lack of knowledge of the media composition to facilitate cultivation of microalgae without the commensal bacteria, the lack of need for high purity cultures for the translational, or the more application-based research that researchers are generally interested.

### 4.4 Potential research gaps

Across the collected literature, many points and inconsistencies in the reportage were noted, that were of significant importance when trying to study and assess axenisation procedures.

#### 4.4.1 Ambiguous glossary

Among the collected literature (including reviews), there is significant ambiguity in the use of terms that essentially allude to various aspects of an organism rather than its culture. Terms such as “culture”, “species”, “strain” and “clonal” have been used interchangeably across the literature although they mean very different things Belcher and Swale (1988). Even within the context of the collated research, the term species refers to the unique taxonomic identity of the organism being investigated. The term strain refers to a subpopulation of a species that have a property (or set of properties) of interest; for example, the strain IMET1 of *Nannochloropsis oceanica* is capable of higher lipid production although it is taxonomically identical to other populations of *N. oceanica* (Y. Ma et al., 2014). Clonal refers to the population that has been derived from a single microalgal cell through asexual reproduction; sharing identical genetic copies and expression. These three terms therefore refer to the investigated organism itself, but not the conditions in which it is grown. The term “culture” refers to the enclosed system in which the organism is grown. Belcher and Swale (1988) explains that, “A culture, as applied to algae, refers to the complex of culture medium, the organism(s) growing in the medium, and the vessel enclosing these.” Unfortunately, referral to the culture itself remains inconsistent and synecdochical, with terms such as “unialgal”, “single-strain”, and “single-culture” used synonymously to imply axeny. These terms may refer to the fact that only one algal species is grown in the culture, but does not necessarily imply the absence of non-algal entities (e.g. bacteria). Thus, inconsistent usage of terms across the discipline contributes to the uncertainty and inaccuracy in the interpretation of the textual reports.

#### 4.4.2 Lack of attention to the medium

The growth medium conditions are inherently different from the natural conditions from where the microalgae samples were collected and consequently, the growth dynamics of the microalgae and the contaminating microbes is expected to change. This is particularly true when employing nutrient-rich heterogenous medium such as Lysogeny Broth which provides a carbon source, and tends to favour bacterial growth over microalgae. Conversely, carbon-deficient medium such as the f medium relies on the ability of the microalgae to use photosynthesis to assimilate inorganic carbon and achieve dominance over heterotrophic bacteria. Corresponding increase in metabolic stress when using chemical methods such as hypochlorite or antibiotics during axenisation process, however, could significantly affect microalgal cell viability and decrease the chances of successful subculturing. This can be demonstrated using the example of *Euglena gracilis*, where, in all three reported cases, high antibiotic concentrations were used to achieve axenicity. However, the two studies that successfully established axenic cultures; namely by Gumińska et al. (2018) and Pappas and Hoffman (1952) employed Cramer-Myers’s medium and Hall’s medium respectively. Both these media recipes include a carbon source to allow microalgal cells to tap into their mixotrophic pathways. In contrast, Jones et al. (1973), who used artificial seawater, found that the microalgae became unviable before the bacteria were affected. Antibiotics caused chloroplast bleaching in Euglena, thereby impairing their ability to fix carbon. The supplementation of a carbon source, as provided by the media used in Gumińska et al. (2018) and Pappas and Hoffman (1952) may have mitigated the antibiotics’ deleterious effects.

Even the manner in which a method is applied in tandem with the medium can affect outcomes. For example, if anoxy is achieved through flushing the medium with sterile CO_2_ (as described by M. Ma et al. (2017)), it can begin to alter the pH of the culture medium. In such instances, consideration to the buffering capacity, for example through the use of TAP medium has a significantly higher buffering capacity owing to the presence of phophate buffer salts compared to BBM, and correspondingly, the effectiveness of the applied technique can vary significantly.

#### 4.4.3 Lack of identification of co-growers

Across the 63 publications only three supplied sufficient information about the contaminating microbes. This is particularly relevant in cases where chemical methods such as phenol or antibiotics were used to achieve axeny as the effectiveness of the applied method is contingent on the relative susceptibility of the contaminating microbe to the treatment in comparison to the microalgae. In other words, if the contaminating / co-growing microbes are more resilient to the applied treatment in comparison to the microalgae, subsequent efforts to reproduce the outcome may fail even if the described procedure is diligently followed. This is particularly relevant in the case of diatoms as demonstrated by Grossart (1999) that even within the same culture different micro-communities of microalgae and bacteria can exist.

#### 4.4.4 Lack of diversity in reportage

A wider collection of literature helps ensure a strong representation of techniques, especially across different lab settings. The 81 diatom species noted in Figure 3 was obtained from just 11 publications in which 11 methods were described to achieve axenic cultures. Likewise for dinoflagellates, the 36 species were noted in 15 publications and described 13 methods that were used to achieve axenic cultures, while for green algae, 31 species subjected to 19 different methods were described in 32 publications. Finally in the matter of microalgal divisions represented in the literature, cryptomonads, eugalenoids, red algae, etc. that were grouped as “other eukaryotes” in Figure 3d, were poorly represented in the collected literature and indicates a potential direction for further research work. A more focused approach towards testing novel methods that is subsequently corroborated by other worker groups / laboratories would significantly increase confidence in the success rates noted for each method. Moreover, since each method (or its combination) has its intrinsic false positive rates, using the same workflow over multiple species, even within the same Division can affect subsequent efforts to reproduce the results. More work is also required on systematic sampling, identification and purification of microalgae from natural sources. Although axenisation research has been carried out in many countries (Table 1), on-field sampling was described in only 19 publications (Figure 2). Type-sampling, archiving and perseveration of microalgae from natural water sources has become increasingly relevant as their local populations have become increasingly sparse and the chance of obtaining rarer species has become more precarious owing to the effects of climate change (Gimenez Papiol, 2017).

### 4.5 Limitations of this study

The work presented here attempted to systematically collect and curate relevant publications, and the extract information based upon which cogent axenisation workflows and strategies could be developed. However, there are aspects of this work that require careful interpretation and requires further research to furnish missing pieces of data.

## 5 Conclusion

Promising workflows circumventing the use of antibiotics appears to be filtration ↔ washing ↔ micropicking for diatoms, micropicking ↔ subculturing ↔ flow cytometry for dinoflagellates, and anoxy ↔ photosensitisation ↔ streak plating for green algae. More high quality reports with other methods and microalgal groups are required to estimate success rates and suggest alternate workflow strategies to achieve axenic eukaryotic microalgal cultures. A combination of at least three tests is recommended to verify successful axenisation that could include cell counting (using growth colonies on agar or flow cytometry), sequencing (16S and/or 18S), and microscopy (epifluorescence, light, confocal, etc.)

## Supplementary materials

**Supplementary Text 1:** Details on data collection and synthesis when assessing each publication.

**Supplementary Text 2:** Details of the questions used to assess the evidence quality of each publication.

**Supplementary awk code:** The script was used to further filter irrelevant titles recovered from database search. This is available in Supplementary Materials, Supplementary Text, Section 2.

**Supplementary Table 1:** Table detailing the parameters, examples, and the rationale of each exclusion criteria. It is available in the document Supplementary materials, Section 3.

**Supplementary Table 2:** Master table of all the data extracted from the publications upon using which network component analyses were performed. This is available as Supplementary_table2.csv.

**Supplementary Table 3:** Species levels counts of positive or negative axenicity outcomes for each method described in literature.

**Supplementary Table 4:** Number of species currently held in four major culture collections.

**Supplementary Figure 1:** Quality score distribution and outcome share of the six most commonly used methods used in experimental workflows. Data presented is at a species level of resolution.

**Supplementary Figure 2:** Overview of species and axenization workflows for other eukaryotic microalgal groups noted from literature.

**Supplementary Figure 3:** Quality score profile of publications across the timeline.

**Supplementary codes:** This is available in file named Supplementary_codes.pdf, It contains the codes and methods used to process the data extracted from literature.

## Supporting information

Supplementary codes

## Acknowledgements

The authors are very thankful to the respondents from the culture banks who took the time to respond to our query. A.I is a grateful recipient of the Irish Research Council (IRC) Postdoctoral Fellowship (GOIPD/2023/1240), and UCD College of Engineering & Architecture Seed Funding Programme 2023 (Career Development Award).

M.M and T.Q are thankful for the Erasmus mobility award to facilitate their training at UCD.

## References

Aray-Andrade, M. M., Uyaguari-Diaz, M. I., & Bermúdez, J. R. (2018). Short-term deleterious effects of standard isolation and cultivation methods on new tropical freshwater microalgae strains. PeerJ, 6, e5143. 10.7717/peerj.5143

Belcher, H., & Swale, E. (1988). Culturing algae: A guide for schools and colleges. Institute of Terrestrial Ecology.

Brandt, J., & Albertsen, M. (2018). Investigation of Detection Limits and the Influence of DNA Extraction and Primer Choice on the Observed Microbial Communities in Drinking Water Samples Using 16S rRNA Gene Amplicon Sequencing. Frontiers in Microbiology, 9, 2140. 10.3389/fmicb.2018.02140

Cappuccino, J. G., & Sherman, N. (2008). Microbiology: A laboratory manual (8th ed). Pearson/Benjamin Cummings.

Chamberlain, S. A., & Szöcs, E. (2013). Taxize: Taxonomic search and retrieval in R. F1000Research, 2, 191. 10.12688/f1000research.2-191.v2

Clauset, A., Newman, M. E. J., & Moore, C. (2004). Finding community structure in very large networks. Physical Review E, 70(6), 066111. 10.1103/PhysRevE.70.066111

Csárdi, G., Nepusz, T., Müller, K., Horvát, S., Traag, V., Zanini, F., & Noom, D. (2024, October). Igraph for R: R interface of the igraph library for graph theory and network analysis. 10.5281/ZENODO.7682609

Deng, Y., Yu, R., Grabe, V., Sommermann, T., Werner, M., Vallet, M., Zerfaß, C., Werz, O., & Pohnert, G. (2024). Bacteria modulate microalgal aging physiology through the induction of extracellular vesicle production to remove harmful metabolites. Nature Microbiology, 9(9), 2356–2368. 10.1038/s41564-024-01746-2

Dextro, R. B., Andreote, A. P., Vaz, M. G., Carvalho, C. R., & Fiore, M. F. (2024). The pros and cons of axenic cultures in cyanobacterial research. Algal Research, 78, 103415. 10.1016/j.algal.2024.103415

Eppstein, D., Löffler, M., & Strash, D. (2010, June). Listing All Maximal Cliques in Sparse Graphs in Near-optimal Time. 10.48550/arXiv.1006.5440

Fernandez-Valenzuela, S., Chávez-Ruvalcaba, F., Beltran-Rocha, J. C., San Claudio, P. M., & Reyna-Martínez, R. (2021). Isolation and Culturing Axenic Microalgae: Mini–Review. The Open Microbiology Journal, 15(1), 111–119. 10.2174/1874285802115010111

Feuersänger, C. (2023). Pgfplots – A TeX Package to Create Normal and Logarithmic Plots in Two and Three Dimensions.

Foster, Z. S. L., Sharpton, T. J., & Grünwald, N. J. (2017). Metacoder: An R package for visualization and manipulation of community taxonomic diversity data (T. Poisot, Ed.). PLOS Computational Biology, 13(2), e1005404. 10.1371/journal.pcbi.1005404

Fouilland, E., Galès, A., Beaugelin, I., Lanouguère, E., Pringault, O., & Leboulanger, C. (2018). Influence of bacteria on the response of microalgae to contaminant mixtures. Chemosphere, 211, 449–455. 10.1016/j.chemosphere.2018.07.161

Francoeur, S. N., Espeland, E. M., & Wetzel, R. G. (2003). Short-term effects of nitrogen and extracellular protease amendment on algal productivity in nitrogen-deprived periphyton. Journal of Freshwater Ecology, 18(1), 105–113. 10.1080/02705060.2003.9663956

Gimenez Papiol, G. (2017). Climate conditions, and changes, affect microalgae communities… should we worry? Integrated Environmental Assessment and Management, 14(2), 181–184. 10.1002/ieam.2009

Grossart, H. (1999). Interactions between marine bacteria and axenic diatoms (Cylindrotheca fusiformis, Nitzschia laevis, and Thalassiosira weissflogii) incubated under various conditions in the lab. Aquatic Microbial Ecology, 19, 1–11. 10.3354/ame019001

Gumińska, N., Płecha, M., Walkiewicz, H., Hałakuc, P., Zakryś, B., & Milanowski, R. (2018). Culture purification and DNA extraction procedures suitable for next-generation sequencing of euglenids. Journal of Applied Phycology, 30(6), 3541–3549. 10.1007/s10811-018-1496-0

Han, J., Wang, S., Zhang, L., Yang, G., Zhao, L., & Pan, K. (2016). A method of batch-purifying microalgae with multiple antibiotics at extremely high concentrations. Chinese Journal of Oceanology and Limnology, 34(1), 79–85. 10.1007/s00343-015-4288-2

Imai, I., & Yamaguchi, M. (1994). A simple technique for establishing axenic cultures of phytoflagellates. Bulletin of Japanese Society of Microbial Ecology, 9(1), 15–17. 10.1264/microbes1986.9.15

Jadad, A. R., Moore, R., Carroll, D., Jenkinson, C., Reynolds, D. M., Gavaghan, D. J., & McQuay, H. J. (1996). Assessing the quality of reports of randomized clinical trials: Is blinding necessary? Controlled Clinical Trials, 17 (1), 1–12. 10.1016/0197-2456(95)00134-4

Jones, A., Rhodes, M. E., & Evans, S. C. (1973). The use of antibiotics to obtain axenic cultures of algae. British Phycological Journal, 8(2), 185–196. 10.1080/00071617300650211

Kan, Y., & Pan, J. (2010). A one-shot solution to bacterial and fungal contamination in the green alga chlamy-domonas reinhardtii culture by using an antibiotic cocktail ^1^. Journal of Phycology, 46(6), 1356–1358. 10.1111/j.1529-8817.2010.00904.x

Lamport, L. (1994). LATEX: A document preparation system: User’s guide and reference manual (2nd ed). Addison-Wesley Pub. Co.

Li, Y., Chen, X., Wang, Q., Liu, Y., Li, J., Gong, Q., & Gao, X. (2022). Diversity and dynamics of bacterial communities associated with Haematococcus pluvialis at different life stages. Journal of Applied Phycology, 34(3), 1353–1361. 10.1007/s10811-022-02729-8

Ma, M., Yuan, D., He, Y., Park, M., Gong, Y., & Hu, Q. (2017). Effective control of Poterioochromonas malhamensis in pilot-scale culture of Chlorella sorokiniana GT-1 by maintaining CO2-mediated low culture pH. Algal Research, 26, 436–444. 10.1016/j.algal.2017.06.023

Ma, Y., Wang, Z., Yu, C., Yin, Y., & Zhou, G. (2014). Evaluation of the potential of 9 Nannochloropsis strains for biodiesel production. Bioresource Technology, 167, 503–509. 10.1016/j.biortech.2014.06.047

McCracken, I. (1969). Purifying algal cultures - a review of chemical methods. Proceedings of the Nova Scotian Institute of Science, 38(1), 145–168.

Molina-Cárdenas, C. A., Sánchez-Saavedra, M. D. P., & Licea-Navarro, A. F. (2016). Decreasing of bacterial content in Isochrysis galbana cultures by using some antibiotics. Revista de biología marina y oceanografía, 51(1), 101–112. 10.4067/S0718-19572016000100010

Pappas, G., & Hoffman, H. (1952). The Use of Antibiotics for Obtaining Bacteria-Free Cultures of Euglena. The Ohio Journal of Science, 52(2), 102–105.

Pendersen, T. L. (2024). Tidygraph: A Tidy API for Graph Manipulation.

Pushpakumara, B. L. D. U., Tandon, K., Willis, A., & Verbruggen, H. (2023). Unravelling microalgal-bacterial interactions in aquatic ecosystems through 16S rRNA gene-based co-occurrence networks. Scientific Reports, 13(1), 2743. 10.1038/s41598-023-27816-9

R Core Team. (2024). R: A Language and Environment for Statistical Computing.

Solomon, W., Mutum, L., Janda, T., & Molnár, Z. (2023). Potential benefit of microalgae and their interaction with bacteria to sustainable crop production. Plant Growth Regulation, 101(1), 53–65. 10.1007/s10725-023-01019-8

Tanau, T. (2021). The TikZ and PGF Packages.

Thain, M., & Hickman, M. (2004). The Penguin dictionary of biology (11th ed). Penguin Books.

Vu, C. H. T., Lee, H.-G., Chang, Y. K., & Oh, H.-M. (2018). Axenic cultures for microalgal biotechnology: Establishment, assessment, maintenance, and applications. Biotechnology Advances, 36(2), 380–396. 10.1016/j.biotechadv.2017.12.018

Wang, L., Yang, F., Chen, H., Fan, Z., Zhou, Y., Lu, J., & Zheng, Y. (2016). Antimicrobial cocktails to control bacterial and fungal contamination in Chlamydomonas reinhardtii cultures. BioTechniques, 60(3), 145–149. 10.2144/000114392

Wickham, H., Averick, M., Bryan, J., Chang, W., McGowan, L., François, R., Grolemund, G., Hayes, A., Henry, L., Hester, J., Kuhn, M., Pedersen, T., Miller, E., Bache, S., Müller, K., Ooms, J., Robinson, D., Seidel, D., Spinu, V., … Yutani, H. (2019). Welcome to the Tidyverse. Journal of Open Source Software, 4(43), 1686. 10.21105/joss.01686

Winter, D. J. (2017). Rentrez: An R package for the NCBI eUtils API The R Journal. The R Journal, 9(2), 520–526.

Youn, J.-Y., & Hur, S.-B. (2007). Antibiotics and their optimum concentration for axenic culture of marine microalgae. ALGAE, 22(3), 229–234. 10.4490/ALGAE.2007.22.3.229

Yu, B. S., Lee, S. Y., & Sim, S. J. (2022). Effective contamination control strategies facilitating axenic cultivation of Haematococcus pluvialis: Risks and challenges. Bioresource Technology, 344, 126289. 10.1016/j.biortech.2021.126289

Yu, G., Smith, D. K., Zhu, H., Guan, Y., & Lam, T. T.-Y. (2017). Ggtree : An r package for visualization and annotation of phylogenetic trees with their covariates and other associated data (G. McInerny, Ed.). Methods in Ecology and Evolution, 8(1), 28–36. 10.1111/2041-210X.12628

