## Supplementary materials for "Using network component analysis to study axenisation strategies for phototrophic eukaryotic microalgae"

#### 1 Supplementary texts

##### 1.1 Biological methods

**1.1.1 Co-culture (CoCu):** Introducing a microbe (typically a bacterium) of known biological characteristics and antibiotic sensitivity into the xenic microalgal culture. This introduced microbe is expected to outgrow or outcompete the contaminating microbes. Subsequently, antibiotics is used to eliminate the introduced microbe, leaving behind an axenic culture.

**1.1.2 Selective Predation (SePd):** Introducing an organism such as an amoeba that engulfs and selectively digests bacteria while sparing microalgae due to their protective cell walls. The amoeba is later removed using selective antibiotics, resulting in an axenic microalgal culture.

**1.1.3 Phototaxis (PhoX):** Leveraging the phototactic movement of motile microalgae, the culture vessel is covered with an opaque material except at the top, where a light source is placed. The phototactic microalgae migrate towards the light, where they can be collected, leaving non-motile contaminating microbes at the bottom.

##### 1.2 Chemical methods

**1.2.1 Antibiotics:** Specific antibiotics, such as streptomycin or kanamycin, are added to the culture to selectively target and eliminate contaminating microbes, yielding an axenic microalgal culture.

**1.2.2 Lysozyme (LysZ):** The enzyme lysozyme is introduced to the culture to degrade bacterial cell walls, thereby reducing bacterial contamination and facilitating the eventual achievement of an axenic culture.

**1.2.3 Hypochlorite (Chlo):** Sodium hypochlorite (NaOCl) is used to treat the culture, causing oxidative stress that inactivates or kills contaminating microbes. The treatment is carefully controlled to preserve the viability of the microalgae.

**1.2.4 Surfactants (Dtrg):** Adding surfactants, such as detergents, disrupts microbial membranes, thereby reducing contaminant loads in the culture and aiding in the establishment of an axenic culture.

**1.2.5 Phenol (Phen):** Phenol and its derivatives are used for their toxic effects on contaminating microbes, allowing microalgal cells to outcompete the contaminants and establish an axenic culture.

**1.2.6 Anoxy (Anox):** Creating an anaerobic environment by sealing the culture vessel to exclude oxygen. Contaminating microbes that require oxygen perish, while microalgae adapted to low-oxygen conditions survive and proliferate.

**1.2.7 Rose Bengal / Photosensitization (PhoS):** The photosensitizing dye Rose Bengal is introduced to the culture and exposed to light. This generates reactive oxygen species that selectively kill contaminating microbes while sparing the microalgae.

**1.2.8 Salt Solution (Salt):** Subjecting the culture to high concentrations of salt (typically, NaCl) creates osmotic stress that kills or inhibits contaminating microbes. Microalgae with high salt tolerance can survive and proliferate.

#### 1.3 Physical methods

**1.3.1 Centrifugation (Ctfg):** The culture is centrifuged at high speeds, causing cells to sediment based on their density. High-density microalgae typically form a pellet at the bottom, which can be collected and processed further to remove contaminants.

**1.3.2 Streak Plating (StPl):** A culture sample is streaked onto an agar medium to isolate individual colonies of microalgae and bacteria. Isolated microalgal colonies can be picked and cultured further to obtain axenic cultures.

**1.3.3 Washing (Wash):** Repeated washing of the culture with sterile buffer or medium removes loosely attached bacteria or spores. This method is often combined with other treatments to achieve axeny.

**1.3.4 Micropicking (Mkpk):** Under a microscope, individual microalgal cells are physically separated from contaminants using fine tools, such as microcapillary. These isolated cells are then cultured in fresh medium.

**1.3.5 Filtration (Fltr):** The culture is passed through a filter with a pore size small enough to retain microalgae while allowing smaller contaminating microbes to pass through. The retained microalgae are then cultured to obtain axenic cultures.

**1.3.6 Density Gradient (DenG):** A density gradient medium, such as Percoll, is used to separate microalgae from contaminants based on differences in buoyant density. The microalgae are collected from the appropriate layer and cultured further.

**1.3.7 Ultrasonication (UltS):** High-frequency sound waves are applied to the culture to disrupt bacterial cells while sparing microalgae. The treated culture can then be subcultured to achieve axenic conditions.

**1.3.8 Subculturing (SubC):** A subsample of the culture, expected to have low contamination levels, is transferred to fresh medium tailored to support microalgae growth (e.g., high salt, no carbon source). This allows microalgae to outgrow contaminants over successive subcultures.

**1.3.9 Microfluidics (Mifl):** Microfluidic devices with thin channels are used to sort and isolate microalgal cells based on size and shape. The isolated cells, presumed free of contaminants, are cultured on selective media to achieve axenic conditions.

**1.3.10 French Press (FrPr):** The culture is passed through a French press, which applies high pressure to disrupt contaminating microbes. Microalgae, which are more resistant to pressure, survive and are subsequently cultured.

**1.3.11 Serial Dilution (SrDl):** A dense microalgal culture is serially diluted to create multiple aliquots. With each dilution, the likelihood of obtaining a contaminant-free sample increases. Axenic cultures can be obtained by culturing aliquots in fresh medium.

###### 1.3.12 Adsorptive Resin (Resn): The culture is passed through a cation exchange resin that binds negatively charged microalgal cells. The adsorbed cells are eluted using a salt solution and cultured to obtain axenic conditions.

#### 1.4 Data synthesis

###### 1.4.1 Publications reporting multiple organisms: In the publications where multiple organisms were reported, only the eukaryotic microalgae were considered and corresponding details were extracted. For example, in (Aray-Andrade et al., 2018), axenisation methods were investigated on *Chlorella pyrenoidosa*, *Chlamydomonas* sp., and *Anacystis nidulans*. Since *A. nidulans* is a prokaryote (heterotypic synonym: *Synechococcus elongatus*, Taxon ID: 1140), it was not included in the final database and only details of *Chlorella pyrenoidosa* and *Chlamydomonas* sp. were collected and summarised.

###### 1.4.2 Data resolution: The publication year, axeny verification/screening method, research quality score, geolocation, and growth media used were collected at the publication level. Data points pertaining to the microalgal species, axenisation workflow and final axeny outcome were collected and summarised at the species level.

#### 2 AWK FILTRATION CODE

The following awk code was used to quickly filter out irrelevant titles:

```
#!/bin/awk -f
BEGIN{ FS=","}
FNR==1 { print $0 > "terms.csv"              # write header row
       print $0 > "notterms.csv"             # to both files
       next
      }
      { name=$2
       sub(/^[ ]+|[ ]+$/,"",name)            # strip leading/trailing spaces
          from name
       if (name~/seaweed/ ||
        name~/biogas/ ||
        name~/bloom/ ||
        name~/porphyra/ ||
        name~/cyano*/ ||
        name~/waste/ ||
        name~/preservation/||
        name~/blue/||
        name~/sludge/ ||
        name~/fish/ ||
        name~/toxic/ ||
        name~/rotifer/ ||
        name~/Schizochytrium/||
        name~/Crypthecodinium/ ||
        name~/Pseudopfiesteria/ ||
        name~/Paramecium/ ||
        name~/raceway/ ||
        name~/aquaculture/ ||
        name~/salmon/)
         print $0 > "terms.csv"
       else
         print $0 > "notterms.csv"
      }
END{}
```

The code generates a workfile terms.csv which contains titles with the irrelevant terms, and another workfile notterms.csv which contains the cleaned list of titles.

### 3 Supplementary Figures

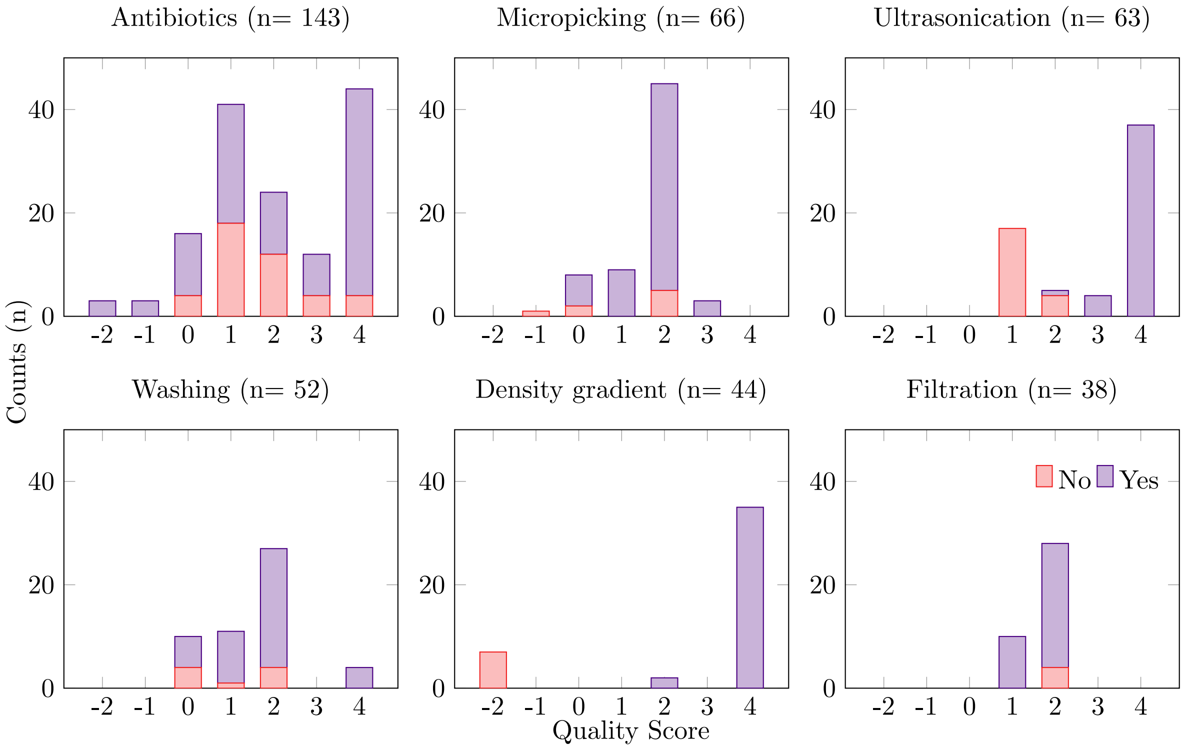


Figure S1: Quality score distribution and outcome share of the six most commonly used methods used in experimental workflows. Data presented is at a species level of resolution.

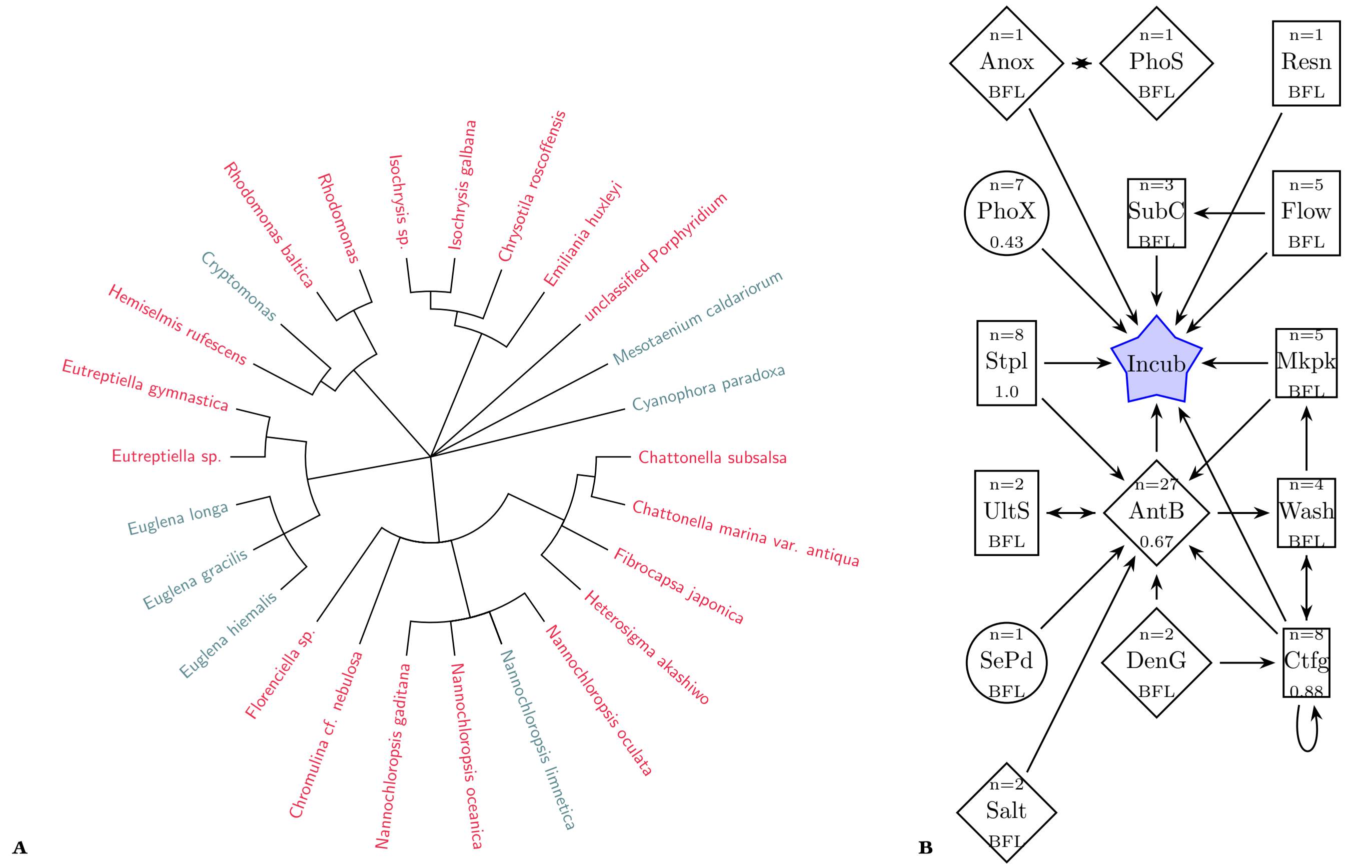


Figure S2: Overview of species and axenization workflows for other eukaryotic microalgal groups noted from literature. (A) Taxonomic tree of 26 microalgal algae species identified across 22 publications. Species names are color-coded: red for marine, sunset-yellow for brackish, and steelblue for freshwater species. (B) Network workflow of 15 methods used to achieve axenic microalgal cultures. Numbers in the network components represent success rates when n>6; for n≤6, success rates are below the filter limit (BFL) and not calculated. Node shapes denote method types: squares for “Physical”, circles for “Biological”, and diamonds for “Chemical”.

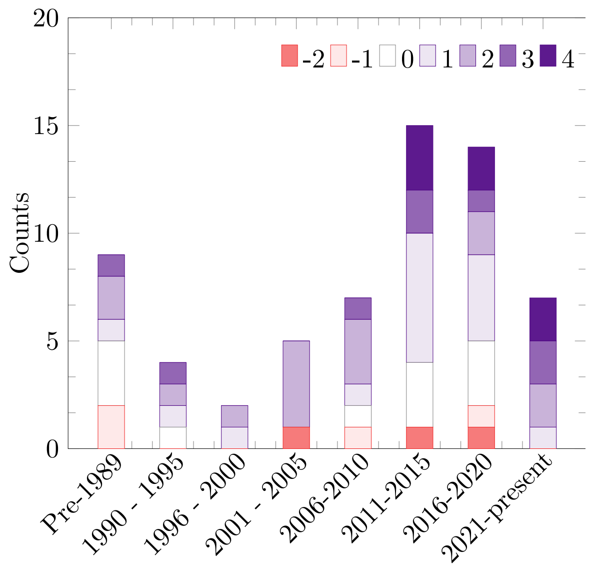


Figure S3: Quality score profile of publications across the timeline.

### 4 Supplementary tables

Table S1: Exclusion Criteria and Reasons

| Parameter | Exclusions | Reason |
|---|---|---|
| Cellular organisation | Multicellular macroalgae were excluded. Studies on marine viruses were also excluded. | Review focuses on unicellular microalgae. |
| Taxonomy | Non-eukaryotic organisms were excluded from the search results. | Review focuses on eukaryotic microalgae. |
| Trophic capability | Heterotrophic microalgae such as *Crypthecodinium cohnii* or *Paramecium caudatum* were excluded. Transient ability to photosynthesise through ingestion of phototrophic microalgae, as in the case of *Pseudopfiesteria shumwayae*, was also excluded. | Review focuses on phototrophic microalgae. |
| Multialgal systems | Publications exclusively focusing on the separation of microalgal species from a co-culture, but without emphasis or details on axeny were excluded. | Separation of microalgal species from a mixture does not imply axeny and without sufficient details, it was impossible to ascertain the status from the literature. |
| Non-experimental literature* | Review articles, book chapters, instruction manuals, posters, patents, presentations, correspondences, etc. were excluded. | The review sought to compare results from experimental procedures. |
| Unreported results | Some experimental reports, comments pieces, and letters-to-editor describe methods used to achieve axeny, but did not present any results. Such literature was also excluded. | Lack of experimental details did not allow for comparison of methods and results across the literature. |
| Tolerance and metabolism | Works describing the use of antibiotics, or any physico-chemical methods to study physiological changes in microalgae were excluded. For example: dose tolerance of microalgae to certain antibiotics, or ability of microalgae to metabolise antibiotics/antimicrobial compounds. | Metabolic capabilities of the microalgae was not the focus of the work, but rather the ability of a chosen method to generate axenic cultures. |
| Cultivation style | Publications describing cultivation systems exposed to open air such as raceways ponds, aeration tanks, etc., were excluded. | Open cultivation styles cannot maintain the sterile conditions. |
| Unclear identity | Publications that detail the axenisation procedure from natural samples, but do not specify/identify the purified microalgae were excluded. | Context for the applied method was required as response to treatment was expected to vary across microalgal species. |
| Artificial contamination | Publications where a specific contaminant was introduced in an axenic microalgal culture to investigate experimental procedures were excluded. | The review sought to help achieve axeny in natural or cultured samples. Introduced contaminants may not truly represent the contaminants generally found in nature. |
| Uncertain classification | Work describing organisms with unclear designation as an algae, such as *Schizochytrium* spp., Rotifers, etc. were excluded. | Analyses was to be restricted to organisms with widely accepted nomenclature and categorisation. |

Table S2: Master table with information from 63 publications collected and summarised.

| Document is available in the file Supplementary Table 2.csv |
|---|

Table S3: Axenisation methods and success rates

| Method | Success | 4 | 3 | 2 | 1 | 0 | -2 | -1 | Total |
|---|---|---|---|---|---|---|---|---|---|
| Anoxy | Yes | 4 | 0 | 0 | 1 | 0 | 0 | 0 | 5 |
| | No | 0 | 0 | 0 | 0 | 0 | 0 | 0 | 0 |
| Photosensitisation | Yes | 4 | 0 | 0 | 0 | 0 | 0 | 0 | 4 |
| | No | 0 | 0 | 0 | 0 | 0 | 0 | 0 | 0 |
| Antibiotics | Yes | 40 | 8 | 12 | 23 | 12 | 3 | 3 | 101 |
| | No | 4 | 4 | 12 | 18 | 4 | 0 | 0 | 42 |
| Streak plating | Yes | 2 | 4 | 3 | 10 | 1 | 0 | 0 | 20 |
| | No | 0 | 0 | 0 | 0 | 2 | 0 | 1 | 3 |
| Phototaxis | Yes | 0 | 1 | 2 | 6 | 1 | 0 | 0 | 10 |
| | No | 0 | 0 | 0 | 7 | 0 | 0 | 3 | 10 |
| Centrifugation | Yes | 4 | 4 | 1 | 3 | 6 | 0 | 0 | 18 |
| | No | 0 | 0 | 1 | 1 | 2 | 10 | 0 | 14 |
| Micropicking | Yes | 0 | 3 | 40 | 9 | 6 | 0 | 0 | 58 |
| | No | 0 | 0 | 5 | 0 | 2 | 0 | 1 | 8 |
| Selective predation | Yes | 0 | 0 | 0 | 0 | 6 | 0 | 0 | 6 |
| | No | 0 | 0 | 0 | 0 | 0 | 0 | 0 | 0 |
| Washing | Yes | 4 | 0 | 23 | 10 | 6 | 0 | 0 | 43 |
| | No | 0 | 0 | 4 | 1 | 4 | 0 | 0 | 9 |
| Filtration | Yes | 0 | 0 | 24 | 10 | 0 | 0 | 0 | 34 |
| | No | 0 | 0 | 4 | 0 | 0 | 0 | 0 | 4 |
| Serial dilution | Yes | 1 | 0 | 3 | 0 | 0 | 0 | 0 | 4 |
| | No | 0 | 0 | 0 | 0 | 0 | 0 | 0 | 0 |
| Co culture | No | 0 | 0 | 0 | 16 | 0 | 0 | 0 | 16 |
| Microfluidics | Yes | 0 | 0 | 0 | 1 | 2 | 0 | 0 | 3 |
| | No | 0 | 0 | 0 | 0 | 0 | 0 | 0 | 0 |
| Triiodide Resin | Yes | 0 | 0 | 0 | 1 | 0 | 0 | 0 | 1 |
| | No | 0 | 0 | 0 | 0 | 0 | 0 | 0 | 0 |

Table S4: Microalgal Collection Data

| As on | Collection | Xenic | Axenic |
|---|---|---|---|
| 24 April 2024 | CCAP, UK | 2717 | 296 |
| 29 May 2024 | Bigelow, USA | 507 | 2209* |
| 26 April 2024 | CSIRO, Australia | 359 (+350 cyanobacteria) | – |
| 30 May 2024 | NIES, Japan | – | 964 |

* Value obtained by subtracting total collection size of 2716.
– Marks info not available.
