## Supplementary codes for "Using network component analysis to study axenisation strategies for phototrophic eukaryotic microalgae"

Ajay Iyer[1,2], Matthias Monissen[3], Qi Teo[1,2], Oskar Modin[4], and Ronald Halim[1,2]

[1]Conway Institute of Biomolecular and Biomedical Research, University College Dublin (UCD) Belfield, Dublin 4, Ireland.
[2]University College Dublin, School of Biosystems and Food Engineering, Belfield, Dublin 4, Ireland.
[3]RWTH Aachen University, Templergraben 55, 52056 Aachen, Germany.
[4]Architecture and Civil Engineering, Chalmers University of Technology, Gothenburg, Sweden.

2025-03-30

#### Load the required libraries

These are the packages that will be used across all the analyses described here.

```
library(tidyverse) #Set of packages for data wrangling and visualisation.
library(formatR) #Package to make this Markdown render correctly.
library(igraph)
library(ggraph)
library(tidygraph)
```

#### Defining useful functions

Two functions are defined here. The first will count the number of times each method is being reported for a given species. The second will break the workflow into contiguous pairs.

**Code chunk 1.1:**

```
get_treatment_counts <- function(treatment_string) {
  treatments <- str_trim(unlist(str_split(treatment_string, "\\>")))
  treatment_counts <- as.data.frame(table(treatments))
  names(treatment_counts) <- c("Treatment", "Count")
  return(treatment_counts)
}
```

**Code chunk 1.2:**

```
process_row <- function(row) {
  no <- row[["No."]]
  treatments <- unlist(strsplit(row[["Workflow"]], " > "))
  outcome <- row[["Outcome"]]
```

```r
  if (length(treatments) < 2) {
    return(data.frame(No. = no, From = treatments, To = NA, Outcome = outcome))
  }

  from <- head(treatments, -1)
  to <- tail(treatments, -1)

  data.frame(No. = no, From = from, To = to, Outcome = outcome)
}
```

### Notes on the raw data

The raw data is available as `.csv` files. The file numbering corresponding the to the numbering scheme in the Supplementary document attached to the main manuscript. Some tables are within the Supplementary document, while others are provided as `.csvs` with their captions and tables noted in the Supplementary document for ease of reference. The same naming scheme will be used in this document as well.

### 1 Workflows

**Code chunk 2:** Import data using read.csv command

#### 1.1 Working with Diatoms

**Code chunk 3:** Filtering the diatoms from the dataset.

```r
Diatoms <- data %>%
    select(c("Division", "Species", "Method.used", "Axeny.achieved",
        "Habitat")) %>%
    filter(Division == "diatoms") %>%
    mutate(No. = 1:nrow(.)) %>%
    rename(Workflow = "Method.used", Outcome = "Axeny.achieved")

summary(Diatoms)
```

```
##    Division           Species            Workflow           Outcome         
##  Length:102         Length:102         Length:102         Length:102        
##  Class :character   Class :character   Class :character   Class :character  
##  Mode  :character   Mode  :character   Mode  :character   Mode  :character  
##                                                                             
##                                                                             
##                                                                             
##    Habitat               No.        
##  Length:102         Min.   :  1.00  
##  Class :character   1st Qu.: 26.25  
##  Mode  :character   Median : 51.50  
##                     Mean   : 51.50  
##                     3rd Qu.: 76.75  
##                     Max.   :102.00  
```

**Code chunk 4:** Calculating the success rates for each method used for diatoms

```
Method.counts <- Diatoms %>%
    rowwise() %>%
    mutate(TreatmentCounts = list(get_treatment_counts(Workflow))) %>%
    unnest(TreatmentCounts, names_repair = "universal") %>%
    group_by(Treatment) %>%
    summarise(Yes_Count = sum(Outcome == "Yes"), Total_Count = n(),
        Yes_Rate = round(Yes_Count/Total_Count, 2))

Method.counts
```

```
## # A tibble: 12 x 4
##    Treatment        Yes_Count Total_Count Yes_Rate
##    <fct>                <int>       <int>    <dbl>
##  1 Antibiotics             52          69     0.75
##  2 Incubation              74         102     0.73
##  3 Filtration              31          35     0.89
##  4 Surfactants              2           2     1
##  5 Washing                 35          39     0.9
##  6 Density gradient        34          41     0.83
##  7 Ultrasonication         37          54     0.69
##  8 Subculturing             1           1     1
##  9 Centrifugation           6          13     0.46
## 10 Micropicking            30          34     0.88
## 11 Co Culture               0          16     0
## 12 Streak plating           1           1     1
```

**Code chunk 5:** Generating the pair-wise methods table for generating the graph.

```
diatom.graph <- Diatoms %>%
    rowwise() %>%
    do(process_row(.)) %>%
    ungroup() %>%
    select(From, To) %>%
    distinct() %>%
    graph_from_data_frame(directed = TRUE)
```

**Code chunk 6:** Identifying the clusters and cliques for diatoms

```
largest_cliques(diatom.graph)
```

```
## [[1]]
## + 4/12 vertices, named, from c68ba36:
## [1] Ultrasonication  Antibiotics      Centrifugation   Density gradient
##
## [[2]]
## + 4/12 vertices, named, from c68ba36:
## [1] Antibiotics Filtration  Washing     Surfactants
```

```r
cluster_leading_eigen(diatom.graph)
```

```
## IGRAPH clustering leading eigenvector, groups: 3, mod: 0.26
## + groups:
##   $`1`
##   [1] "Antibiotics"    "Co Culture"     "Streak plating" "Incubation"
##
##   $`2`
##   [1] "Filtration"   "Surfactants"  "Washing"      "Subculturing" "Micropicking"
##
##   $`3`
##   [1] "Ultrasonication"  "Density gradient" "Centrifugation"
##
```

```r
median(degree(diatom.graph, loops = TRUE, normalized = FALSE))
```

```
## [1] 4
```

```r
edge_density(diatom.graph, loops = TRUE)
```

```
## [1] 0.2152778
```

**Code chunk 7:** Subgraphs of the clusters for diatoms

```r
diatom.subgraph1 <- induced_subgraph(diatom.graph, c("Antibiotics",
    "Co Culture", "Streak plating"))
edge_density(diatom.subgraph1, loops = TRUE)
```

```
## [1] 0.4444444
```

```r
largest_cliques(diatom.subgraph1)
```

```
## [[1]]
## + 2/3 vertices, named, from 0917891:
## [1] Streak plating Antibiotics
##
## [[2]]
## + 2/3 vertices, named, from 0917891:
## [1] Antibiotics Co Culture
```

```r
median(degree(diatom.subgraph1, loops = TRUE, normalized = FALSE))
```

```
## [1] 2
```

```r
diatom.subgraph2 <- induced_subgraph(diatom.graph, c("Filtration",
    "Surfactants", "Washing", "Subculturing", "Micropicking"))
edge_density(diatom.subgraph2, loops = TRUE)
```

```
## [1] 0.36
```

```
median(degree(diatom.subgraph2, loops = TRUE, normalized = FALSE))
```

```
## [1] 2
```

```
largest_cliques(diatom.subgraph2)
```

```
## [[1]]
## + 3/5 vertices, named, from bccdaeb:
## [1] Filtration  Surfactants Washing
```

```
diatom.subgraph3 <- induced_subgraph(diatom.graph, c("Ultrasonication",
    "Density gradient", "Centrifugation"))
edge_density(diatom.subgraph3, loops = TRUE)
```

```
## [1] 0.4444444
```

```
median(degree(diatom.subgraph3, loops = TRUE, normalized = FALSE))
```

```
## [1] 2
```

```
largest_cliques(diatom.subgraph3)
```

```
## [[1]]
## + 3/3 vertices, named, from d6e6594:
## [1] Ultrasonication  Density gradient Centrifugation
```

**Code chunk 8:** network graph for diatoms

```
as_tbl_graph(diatom.graph) %>% #
  ggraph(layout = "fr")+  # Fruchterman-Reingold layout
  geom_edge_link() +  # Edges
  geom_edge_loop()+ # Loops
  geom_edge_density(fill="blue") + # Density
  geom_node_text(aes(label = name), repel = TRUE, size = 4) + # Labels
  scale_color_viridis_d(option = "C", name = "SCC") +  # Discrete scale for SCCs
  theme_minimal() +
  labs(title = "Diatoms workflow network",
       edge_alpha = "Edge visibility") +
  theme(legend.position = "none",
        axis.title = element_blank(),
        axis.text = element_blank(),
        panel.grid = element_blank())
```

Diatoms workflow network

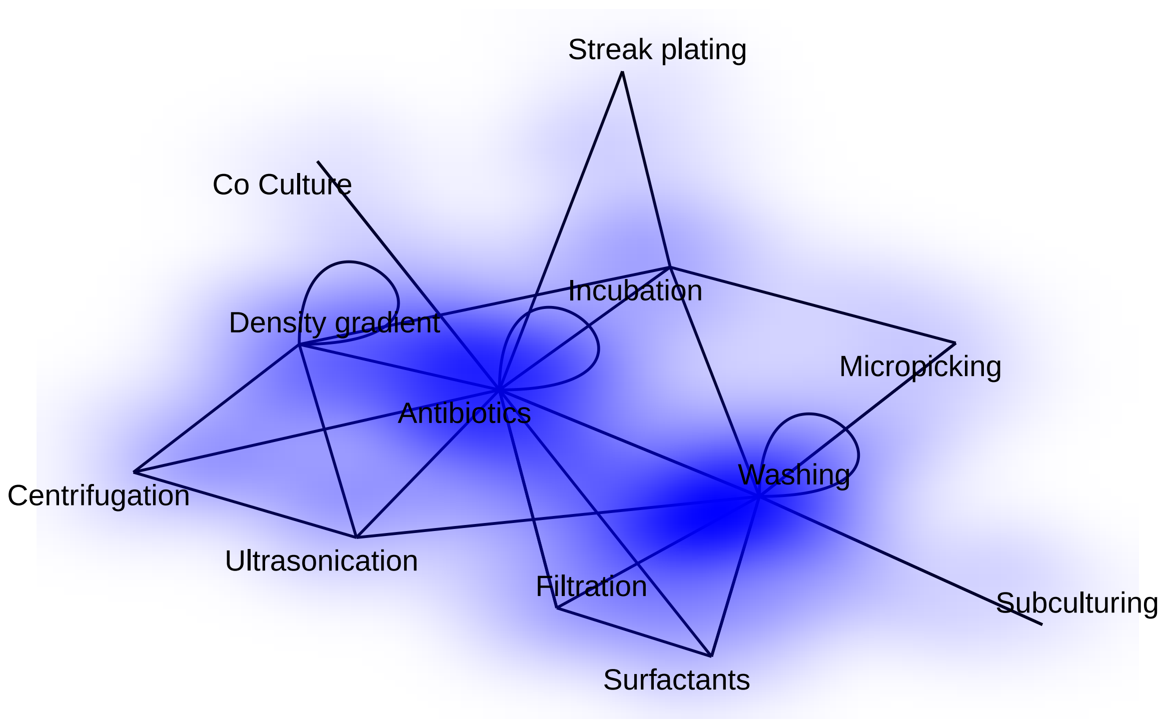


The same graph was reconstructed in LaTeX

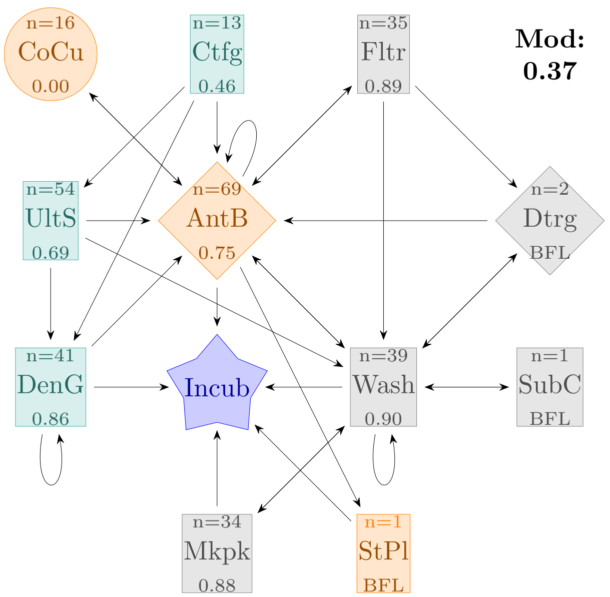


Figure 1: Figure remade using LaTeX. Corresponding Figure 3 in main document

#### 1.2 Working with Dinoflagellates

**Code chunk 9:** Filtering the dinoflagellates from the dataset.

```
Dinoflagellates <- data %>%
    select(c("Division", "Species", "Method.used", "Axeny.achieved",
        "Habitat")) %>%
    filter(Division == "dinoflagellates") %>%
    mutate(No. = 1:nrow(.)) %>%
    rename(Workflow = "Method.used", Outcome = "Axeny.achieved")

summary(Dinoflagellates)
```

```
##    Division           Species            Workflow           Outcome
##  Length:55          Length:55          Length:55          Length:55
##  Class :character   Class :character   Class :character   Class :character
##  Mode  :character   Mode  :character   Mode  :character   Mode  :character
##
##
##
##    Habitat               No.
##  Length:55          Min.   : 1.0
##  Class :character   1st Qu.:14.5
##  Mode  :character   Median :28.0
##                     Mean   :28.0
##                     3rd Qu.:41.5
##                     Max.   :55.0
```

**Code chunk 10:** Calculating the success rates for each method used for dinoflagellates

```
Method.counts <- Dinoflagellates %>%
    rowwise() %>%
    mutate(TreatmentCounts = list(get_treatment_counts(Workflow))) %>%
    unnest(TreatmentCounts, names_repair = "universal") %>%
    group_by(Treatment) %>%
    summarise(Yes_Count = sum(Outcome == "Yes"), Total_Count = n(),
        Yes_Rate = round(Yes_Count/Total_Count, 2))

Method.counts
```

```
## # A tibble: 14 x 4
##    Treatment           Yes_Count Total_Count Yes_Rate
##    <fct>                   <int>       <int>    <dbl>
##  1 Antibiotics                11          18     0.61
##  2 Centrifugation              4           5     0.8
##  3 Incubation                 39          55     0.71
##  4 Micropicking               20          21     0.95
##  5 Selective predation         3           3     1
##  6 Washing                     4           4     1
##  7 Phototaxis                  5          11     0.45
##  8 Filtration                  3           3     1
##  9 Serial dilution             3           3     1
## 10 Lysozyme                    0           1     0
```

```
## 11 Subculturing                5           6     0.83
## 12 Surfactants                 0           1     0
## 13 Flow Cytometry             12          13     0.92
## 14 Density gradient            1           1     1
```

**Code chunk 11:** Generating the pair-wise methods table for generating the graph.

```
dinoflagellate.graph <- Dinoflagellates %>%
    rowwise() %>%
    do(process_row(.)) %>%
    ungroup() %>%
    select(From, To) %>%
    distinct() %>%
    graph_from_data_frame(directed = TRUE)
```

**Code chunk 12:** Identifying the clusters and cliques for dinoflagellates.

```
largest_cliques(dinoflagellate.graph)
```

```
## [[1]]
## + 4/14 vertices, named, from f4aa227:
## [1] Subculturing   Incubation     Flow Cytometry Micropicking
```

```
cluster_leading_eigen(dinoflagellate.graph)
```

```
## IGRAPH clustering leading eigenvector, groups: 4, mod: 0.32
## + groups:
##   $`1`
##   [1] "Selective predation" "Antibiotics"         "Phototaxis"
##   [4] "Filtration"          "Serial dilution"
##
##   $`2`
##   [1] "Micropicking"   "Subculturing"   "Flow Cytometry" "Incubation"
##
##   $`3`
##   [1] "Washing"
##
##   + ... omitted several groups/vertices
```

```
median(degree(dinoflagellate.graph, loops = TRUE, normalized = FALSE))
```

```
## [1] 4.5
```

```
edge_density(dinoflagellate.graph, loops = TRUE)
```

```
## [1] 0.1734694
```

**Code chunk 13:** Subgraphs of the clusters for dinoflagellates

```r
dinoflagellate.subgraph1 <- induced_subgraph(dinoflagellate.graph,
    c("Selective predation", "Antibiotics", "Washing"))
edge_density(dinoflagellate.subgraph1, loops = TRUE)
```

```
## [1] 0.4444444
```

```r
median(degree(dinoflagellate.subgraph1, loops = TRUE, normalized = FALSE))
```

```
## [1] 3
```

```r
largest_cliques(dinoflagellate.subgraph1)
```

```
## [[1]]
## + 2/3 vertices, named, from f428806:
## [1] Selective predation Antibiotics
##
## [[2]]
## + 2/3 vertices, named, from f428806:
## [1] Antibiotics Washing
```

```r
dinoflagellate.subgraph2 <- induced_subgraph(dinoflagellate.graph,
    c("Micropicking", "Subculturing", "Flow Cytometry"))
edge_density(dinoflagellate.subgraph2, loops = TRUE)
```

```
## [1] 0.6666667
```

```r
median(degree(dinoflagellate.subgraph2, loops = TRUE, normalized = FALSE))
```

```
## [1] 4
```

```r
largest_cliques(dinoflagellate.subgraph2)
```

```
## [[1]]
## + 3/3 vertices, named, from e532501:
## [1] Micropicking   Subculturing   Flow Cytometry
```

```r
dinoflagellate.subgraph3 <- induced_subgraph(dinoflagellate.graph,
    c("Centrifugation", "Lysozyme", "Surfactants", "Density gradient"))
edge_density(dinoflagellate.subgraph3, loops = TRUE)
```

```
## [1] 0.3125
```

```r
median(degree(dinoflagellate.subgraph3, loops = TRUE, normalized = FALSE))
```

```
## [1] 2
```

```r
largest_cliques(dinoflagellate.subgraph3)
```

```
## [[1]]
## + 3/4 vertices, named, from 4b074d1:
## [1] Centrifugation Lysozyme       Surfactants
```

```r
dinoflagellate.subgraph4 <- induced_subgraph(dinoflagellate.graph,
    c("Filtration", "Phototaxis", "Serial dilution"))
edge_density(dinoflagellate.subgraph4, loops = TRUE)
```

```
## [1] 0.3333333
```

```r
median(degree(dinoflagellate.subgraph4, loops = TRUE, normalized = FALSE))
```

```
## [1] 2
```

```r
largest_cliques(dinoflagellate.subgraph4)
```

```
## [[1]]
## + 2/3 vertices, named, from bed88a5:
## [1] Serial dilution Phototaxis
##
## [[2]]
## + 2/3 vertices, named, from bed88a5:
## [1] Phototaxis Filtration
```

**Code chunk 14:** Network graph for dinoflagellates

```r
as_tbl_graph(dinoflagellate.graph) %>%
  ggraph(layout = "fr")+  # Fruchterman-Reingold layout
  geom_edge_link() +  # Edges
  geom_edge_loop()+ # Loops
  geom_edge_density(fill="blue") + # Density
  geom_node_text(aes(label = name), repel = TRUE, size = 4) + # Labels
  scale_color_viridis_d(option = "C", name = "SCC") +  # Discrete scale for SCCs
  theme_minimal() +
  labs(title = "Dinoflagellates workflow network",
       edge_alpha = "Edge visibility") +
  theme(legend.position = "none",
        axis.title = element_blank(),
        axis.text = element_blank(),
        panel.grid = element_blank())
```

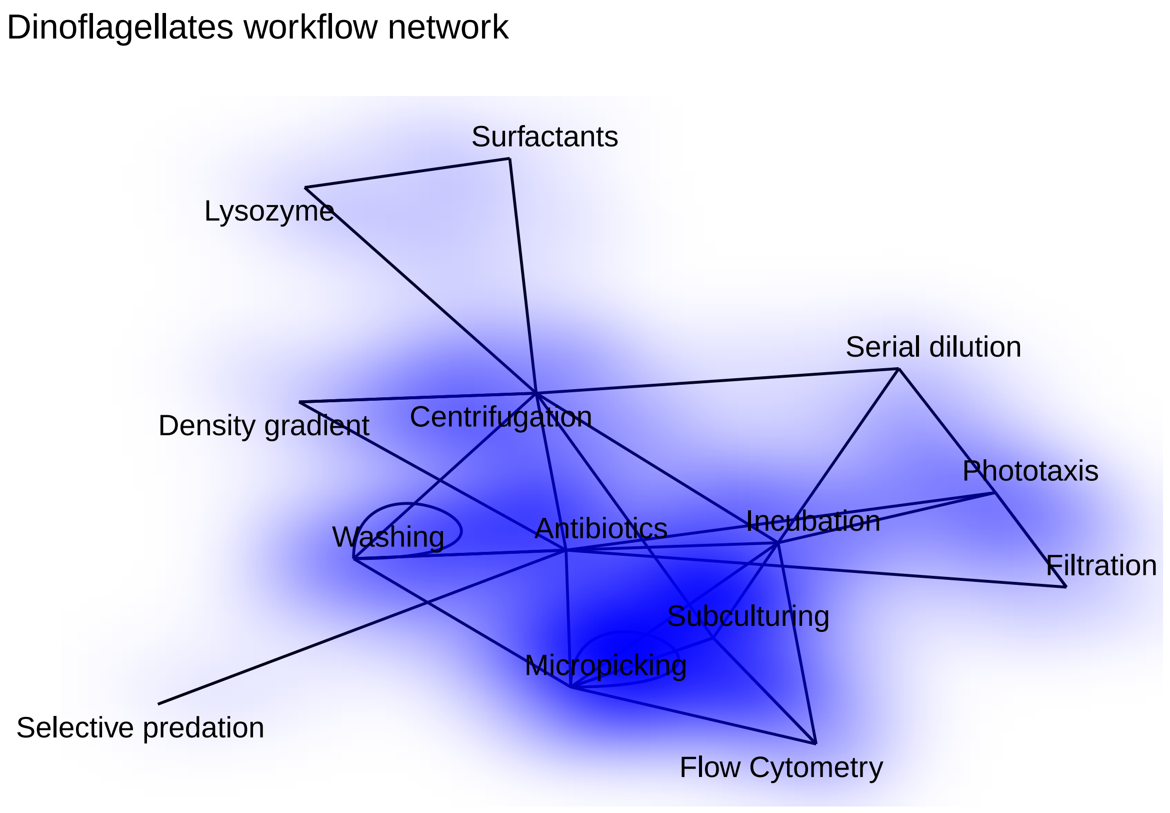


The same graph was reconstructed in LaTeX

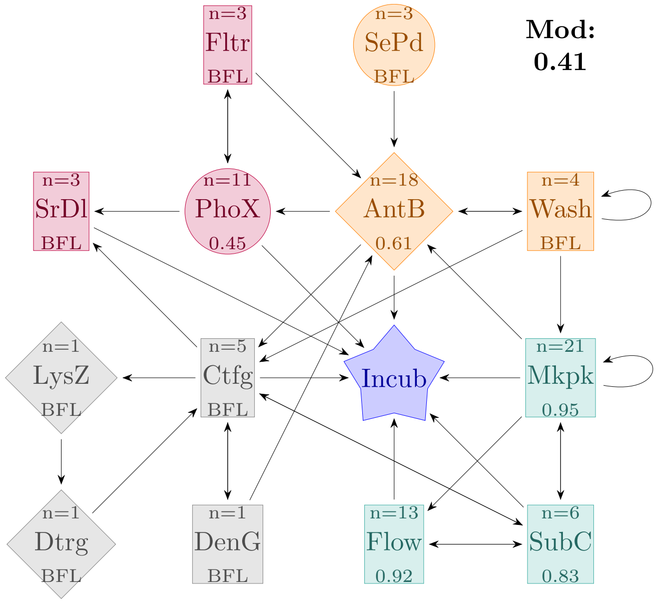


Figure 2: Figure remade using LaTeX. Corresponding Figure 4 in main document

#### 1.3 Working with green algae

**Code chunk 15:** Filtering the green algae from the dataset.

```
Greenalgae <- data %>%
    select(c("Division", "Species", "Method.used", "Axeny.achieved",
        "Habitat")) %>%
    filter(Division == "green algae") %>%
    mutate(No. = 1:nrow(.)) %>%
    rename(Workflow = "Method.used", Outcome = "Axeny.achieved")

summary(Greenalgae)
```

```
##    Division           Species            Workflow           Outcome
##  Length:57          Length:57          Length:57          Length:57
##  Class :character   Class :character   Class :character   Class :character
##  Mode  :character   Mode  :character   Mode  :character   Mode  :character
##
##
##
##    Habitat               No.
##  Length:57          Min.   : 1
##  Class :character   1st Qu.:15
##  Mode  :character   Median :29
##                     Mean   :29
##                     3rd Qu.:43
##                     Max.   :57
```

**Code chunk 16:** Calculating the success rates for each method used for green algae

```
Method.counts <- Greenalgae %>%
    rowwise() %>%
    mutate(TreatmentCounts = list(get_treatment_counts(Workflow))) %>%
    unnest(TreatmentCounts, names_repair = "universal") %>%
    group_by(Treatment) %>%
    summarise(Yes_Count = sum(Outcome == "Yes"), Total_Count = n(),
        Yes_Rate = round(Yes_Count/Total_Count, 2))

Method.counts
```

```
## # A tibble: 20 x 4
##    Treatment          Yes_Count Total_Count Yes_Rate
##    <fct>                  <int>       <int>    <dbl>
##  1 Anoxy                      4           4     1
##  2 Incubation                39          57     0.68
##  3 Photosensitisation         3           3     1
##  4 Antibiotics               20          29     0.69
##  5 Streak plating            11          14     0.79
##  6 Phototaxis                 2           2     1
##  7 Predation                  0           1     0
##  8 Salt solution              1           3     0.33
##  9 Ultrasonication            5           7     0.71
## 10 Flow Cytometry            12          14     0.86
```

```
## 11 Subculturing               1           1     1
## 12 Washing                   1           5     0.2
## 13 Centrifugation            1           6     0.17
## 14 Hypochlorite              0           3     0
## 15 Serial dilution           1           1     1
## 16 Micropicking              3           6     0.5
## 17 Phenol                    0           2     0
## 18 Surfactants               0           2     0
## 19 French Press              0           1     0
## 20 Microfluidics             3           3     1
```

**Code chunk 17:** Generating the pair-wise methods table for generating the graph.

```
greenal.graph <- Greenalgae %>%
    rowwise() %>%
    do(process_row(.)) %>%
    ungroup() %>%
    select(From, To) %>%
    distinct() %>%
    graph_from_data_frame(directed = TRUE)
```

**Code chunk 18:** dentifying the clusters and cliques for green algae

```
largest_cliques(greenal.graph)
```

```
## [[1]]
## + 4/20 vertices, named, from 9f4e7e4:
## [1] Centrifugation Antibiotics    Phototaxis     Streak plating
```

```
cluster_leading_eigen(greenal.graph)
```

```
## IGRAPH clustering leading eigenvector, groups: 5, mod: 0.34
## + groups:
##   $`1`
##   [1] "Anoxy"              "Photosensitisation"
##
##   $`2`
##   [1] "Antibiotics"     "Salt solution"   "Washing"         "Serial dilution"
##
##   $`3`
##   [1] "Streak plating" "Phototaxis"     "French Press"
##
##   $`4`
##   + ... omitted several groups/vertices
```

```
median(degree(greenal.graph, loops = TRUE, normalized = FALSE))
```

```
## [1] 2.5
```

```r
edge_density(greenal.graph, loops = TRUE)
```

```
## [1] 0.1075
```

**Code chunk 19:** Subgraphs of the clusters for green algae

```r
greenal.subgraph1 <- induced_subgraph(greenal.graph, c("Anoxy",
    "Photosensitisation"))
edge_density(greenal.subgraph1, loops = TRUE)
```

```
## [1] 0.5
```

```r
median(degree(greenal.subgraph1, loops = TRUE, normalized = FALSE))
```

```
## [1] 2
```

```r
largest_cliques(greenal.subgraph1)
```

```
## [[1]]
## + 2/2 vertices, named, from 680740a:
## [1] Anoxy              Photosensitisation
```

```r
greenal.subgraph2 <- induced_subgraph(greenal.graph, c("Antibiotics",
    "Salt solution", "Ultrasonication", "Washing", "Centrifugation",
    "Serial dilution"))
edge_density(greenal.subgraph2, loops = TRUE)
```

```
## [1] 0.3333333
```

```r
median(degree(greenal.subgraph2, loops = TRUE, normalized = FALSE))
```

```
## [1] 4
```

```r
largest_cliques(greenal.subgraph2)
```

```
## [[1]]
## + 3/6 vertices, named, from 6fa489e:
## [1] Serial dilution Antibiotics     Washing
##
## [[2]]
## + 3/6 vertices, named, from 6fa489e:
## [1] Antibiotics     Centrifugation  Ultrasonication
##
## [[3]]
## + 3/6 vertices, named, from 6fa489e:
## [1] Antibiotics    Centrifugation Washing
```

```r
greenal.subgraph3 <- induced_subgraph(greenal.graph, c("Streak plating",
    "Phototaxis", "Micropicking", "French Press"))
edge_density(greenal.subgraph3, loops = TRUE)
```

```
## [1] 0.375
```

```r
median(degree(greenal.subgraph3, loops = TRUE, normalized = FALSE))
```

```
## [1] 2.5
```

```r
largest_cliques(greenal.subgraph3)
```

```
## [[1]]
## + 2/4 vertices, named, from 8a6e312:
## [1] Micropicking   Streak plating
##
## [[2]]
## + 2/4 vertices, named, from 8a6e312:
## [1] Phototaxis     Streak plating
##
## [[3]]
## + 2/4 vertices, named, from 8a6e312:
## [1] Streak plating French Press
```

```r
greenal.subgraph4 <- induced_subgraph(greenal.graph, c("Flow Cytometry",
    "Predation", "Subculturing", "Hypochlorite", "Microfluidics"))
edge_density(greenal.subgraph4, loops = TRUE)
```

```
## [1] 0.04
```

```r
median(degree(greenal.subgraph4, loops = TRUE, normalized = FALSE))
```

```
## [1] 0
```

```r
largest_cliques(greenal.subgraph4)
```

```
## [[1]]
## + 2/5 vertices, named, from 885d702:
## [1] Flow Cytometry Subculturing
```

```r
greenal.subgraph5 <- induced_subgraph(greenal.graph, c("Surfactants",
    "Phenol"))
edge_density(greenal.subgraph5, loops = TRUE)
```

```
## [1] 0.25
```

```r
median(degree(greenal.subgraph5, loops = TRUE, normalized = FALSE))
```

```
## [1] 1
```

```r
largest_cliques(greenal.subgraph5)
```

```
## [[1]]
## + 2/2 vertices, named, from 31ef418:
## [1] Surfactants Phenol
```

**Code chunk 20:** Network graph for green algae

```r
as_tbl_graph(greenal.graph) %>%#
  ggraph(layout = "fr")+  # Fruchterman-Reingold layout
  geom_edge_link() +  # Edges
  geom_edge_loop()+ # Loops
  geom_edge_density(fill="blue") + # Density
  geom_node_text(aes(label = name), repel = TRUE, size = 4) + # Labels
  scale_color_viridis_d(option = "C", name = "SCC") +  # Discrete scale for SCCs
  theme_minimal() +
  labs(title = "Green algae workflow network",
       edge_alpha = "Edge visibility") +
  theme(legend.position = "none",
        axis.title = element_blank(),
        axis.text = element_blank(),
        panel.grid = element_blank())
```

Green algae workflow network

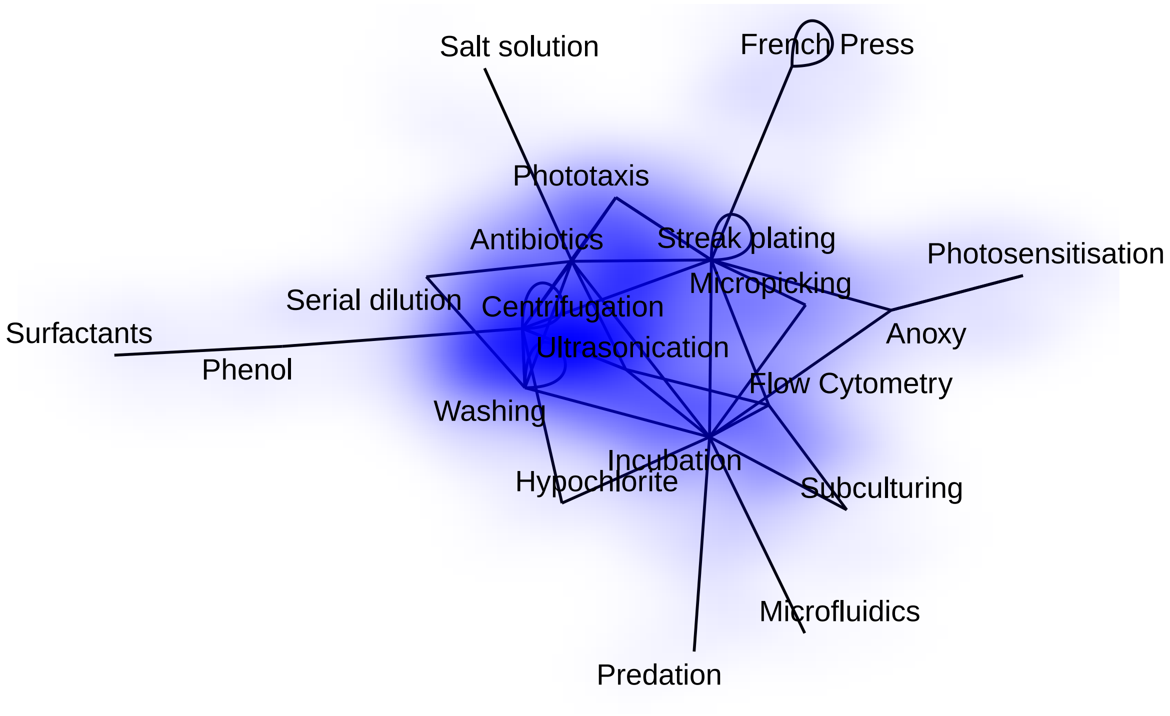


The same graph was reconstructed in LaTeX

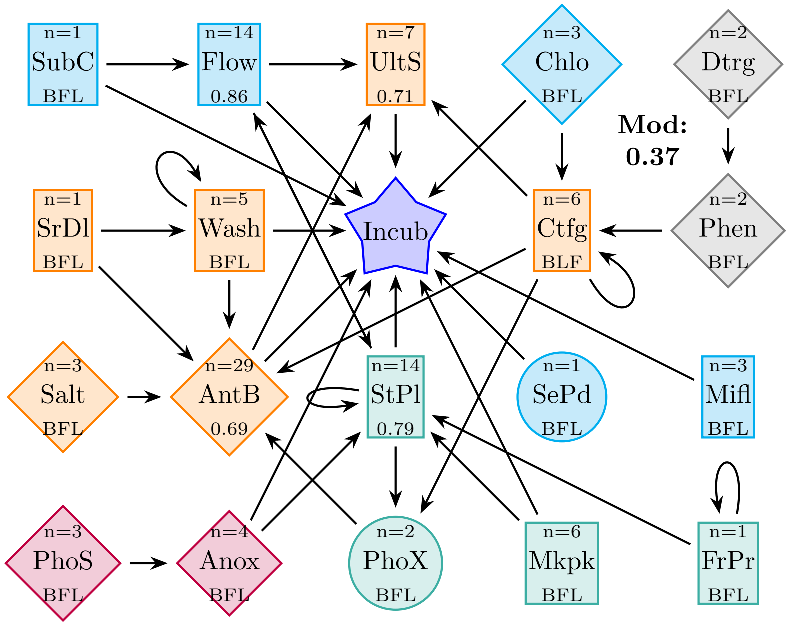


Figure 3: Figure remade using LaTeX. Corresponding Figure 5 in main document

#### 1.4 Working with other eukaryotes

**Code chunk 21:** Filtering the remaining eukaryotic microalgae from the dataset.

```
other <- data %>%
    select(c("Division", "Species", "Method.used", "Axeny.achieved",
        "Habitat")) %>%
    filter(Division != "diatoms" & Division != "green algae" &
        Division != "dinoflagellates") %>%
    mutate(No. = 1:nrow(.)) %>%
    rename(Workflow = "Method.used", Outcome = "Axeny.achieved")

summary(other)
```

```
##    Division           Species            Workflow           Outcome
##  Length:43          Length:43          Length:43          Length:43
##  Class :character   Class :character   Class :character   Class :character
##  Mode  :character   Mode  :character   Mode  :character   Mode  :character
##
##
##
##    Habitat               No.
##  Length:43          Min.   : 1.0
##  Class :character   1st Qu.:11.5
##  Mode  :character   Median :22.0
##                     Mean   :22.0
##                     3rd Qu.:32.5
##                     Max.   :43.0
```

**Code chunk 22:** Calculating the success rates for each method

```
Method.counts <- other %>%
    rowwise() %>%
    mutate(TreatmentCounts = list(get_treatment_counts(Workflow))) %>%
    unnest(TreatmentCounts, names_repair = "universal") %>%
    group_by(Treatment) %>%
    summarise(Yes_Count = sum(Outcome == "Yes"), Total_Count = n(),
        Yes_Rate = round(Yes_Count/Total_Count, 2))

Method.counts
```

```
## # A tibble: 16 x 4
##    Treatment           Yes_Count Total_Count Yes_Rate
##    <fct>                   <int>       <int>    <dbl>
##  1 Anoxy                       1           1     1
##  2 Incubation                 30          43     0.7
##  3 Photosensitisation          1           1     1
##  4 Antibiotics                18          27     0.67
##  5 Phototaxis                  3           7     0.43
##  6 Centrifugation              7           8     0.88
##  7 Micropicking                5           5     1
##  8 Selective predation         3           3     1
##  9 Washing                     3           4     0.75
```

```
## 10 Salt solution               0          2     0
## 11 Ultrasonication             0          2     0
## 12 Flow Cytometry              5          5     1
## 13 Subculturing                3          3     1
## 14 Streak plating              8          8     1
## 15 Density gradient            2          2     1
## 16 Triiodide Resin             1          1     1
```

**Code chunk 23:** Generating the pair-wise methods table for generating the graph.

```
other.graph <- other %>%
    rowwise() %>%
    do(process_row(.)) %>%
    ungroup() %>%
    select(From, To) %>%
    distinct() %>%
    graph_from_data_frame(directed = TRUE)
```

**Code chunk 24:** Network graph for the remaining eukaryotes

```
as_tbl_graph(other.graph) %>%#
  ggraph(layout = "fr")+  # Fruchterman-Reingold layout
  geom_edge_link() +  # Edges
  geom_edge_loop()+ # Loops
  geom_edge_density(fill="blue") + # Density
  geom_node_text(aes(label = name), repel = TRUE, size = 4) + # Labels
  scale_color_viridis_d(option = "C", name = "SCC") +  # Discrete scale for SCCs
  theme_minimal() +
  labs(title = "Other eukaryotes workflow network",
       edge_alpha = "Edge visibility") +
  theme(legend.position = "none",
        axis.title = element_blank(),
        axis.text = element_blank(),
        panel.grid = element_blank())
```

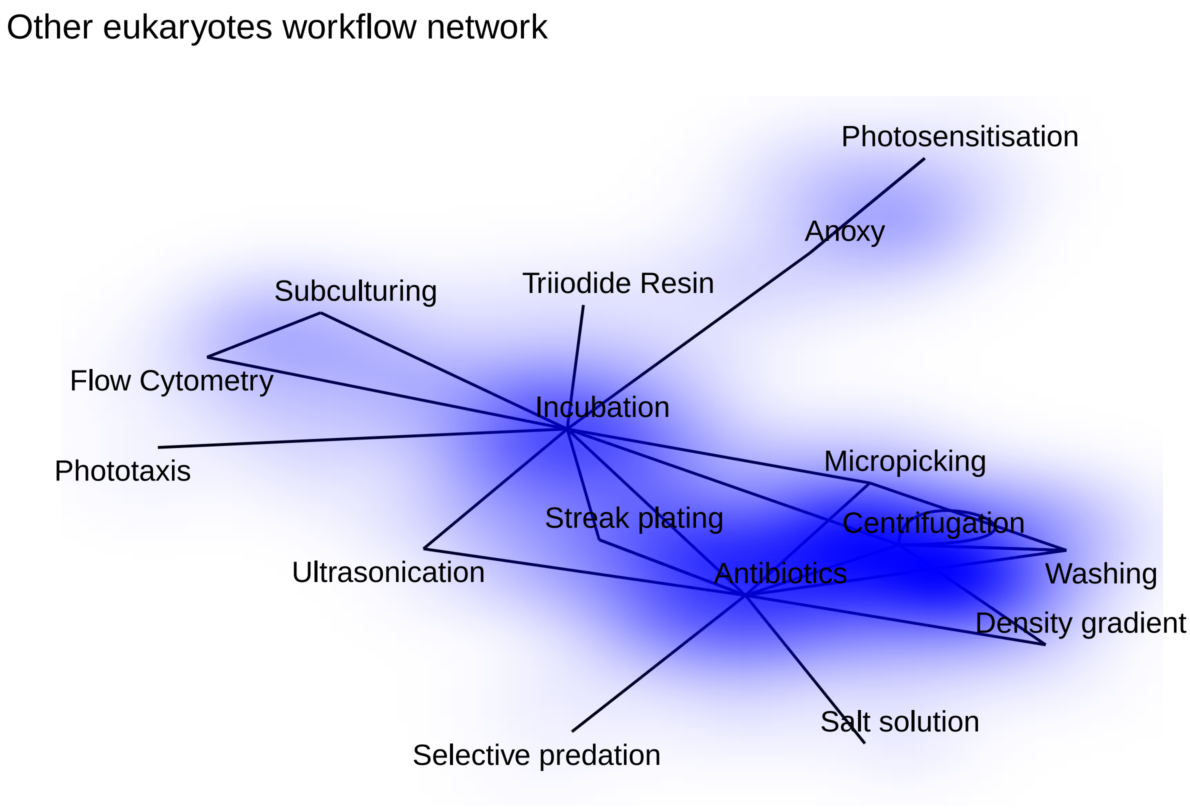


The same graph was reconstructed in LaTeX

```
sessionInfo()
```

```
## R version 4.4.3 (2025-02-28)
## Platform: x86_64-pc-linux-gnu
## Running under: Linux Mint 22.1
##
## Matrix products: default
## BLAS/LAPACK: /usr/lib/x86_64-linux-gnu/openblas-pthread/libopenblasp-r0.3.26.so;  LAPACK version 3.1
##
## locale:
##  [1] LC_CTYPE=en_IE.UTF-8       LC_NUMERIC=C
##  [3] LC_TIME=en_IE.UTF-8        LC_COLLATE=en_IE.UTF-8
##  [5] LC_MONETARY=en_IE.UTF-8    LC_MESSAGES=en_IE.UTF-8
##  [7] LC_PAPER=en_IE.UTF-8       LC_NAME=C
##  [9] LC_ADDRESS=C               LC_TELEPHONE=C
## [11] LC_MEASUREMENT=en_IE.UTF-8 LC_IDENTIFICATION=C
##
## time zone: Europe/Dublin
## tzcode source: system (glibc)
##
## attached base packages:
## [1] stats     graphics  grDevices utils     datasets  methods   base
##
## other attached packages:
```

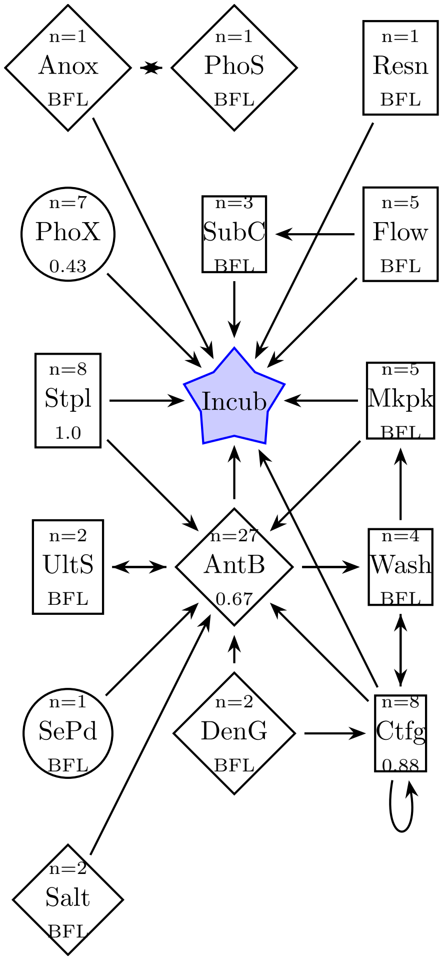


Figure 4: Figure remade using LaTeX. Corresponding Figure 6 in main document

```
##  [1] tidygraph_1.3.1 ggraph_2.2.1    igraph_2.1.4    formatR_1.14
##  [5] lubridate_1.9.4 forcats_1.0.0   stringr_1.5.1   dplyr_1.1.4
##  [9] purrr_1.0.4     readr_2.1.5     tidyr_1.3.1     tibble_3.2.1
## [13] ggplot2_3.5.1   tidyverse_2.0.0
##
## loaded via a namespace (and not attached):
##  [1] viridis_0.6.5      utf8_1.2.4         generics_0.1.3     stringi_1.8.4
##  [5] hms_1.1.3          digest_0.6.37      magrittr_2.0.3     evaluate_1.0.3
##  [9] grid_4.4.3         timechange_0.3.0   fastmap_1.2.0      ggrepel_0.9.6
## [13] gridExtra_2.3      viridisLite_0.4.2  scales_1.3.0       tweenr_2.0.3
## [17] cli_3.6.4          rlang_1.1.5        graphlayouts_1.2.2 polyclip_1.10-7
## [21] munsell_0.5.1      cachem_1.1.0       withr_3.0.2        yaml_2.3.10
## [25] tools_4.4.3        tzdb_0.4.0         memoise_2.0.1      colorspace_2.1-1
## [29] vctrs_0.6.5        R6_2.6.1           lifecycle_1.0.4    MASS_7.3-65
## [33] pkgconfig_2.0.3    pillar_1.10.1      gtable_0.3.6       glue_1.8.0
## [37] Rcpp_1.0.14        ggforce_0.4.2      xfun_0.51          tidyselect_1.2.1
## [41] rstudioapi_0.17.1  knitr_1.49         farver_2.1.2       htmltools_0.5.8.1
## [45] labeling_0.4.3     rmarkdown_2.29     compiler_4.4.3
```
